# Development of novel *in vitro* human alveolar epithelial cell models to study distal lung biology and disease

**DOI:** 10.1101/2020.12.25.424415

**Authors:** Evelyn Tran, Tuo Shi, Xiuwen Li, Adnan Y. Chowdhury, Du Jiang, Yixin Liu, Hongjun Wang, Chunli Yan, William D. Wallace, Rong Lu, Amy L. Ryan, Crystal N. Marconett, Beiyun Zhou, Zea Borok, Ite A. Offringa

**Author notes:** For correspondence: Ite A. Offringa.

## Abstract

Many acute and chronic lung diseases affect the distal lung alveoli. Although airway-derived human cell lines exist, alveolar epithelial cell (AEC)-derived lines are needed to better model these diseases. We have generated and characterized novel immortalized cell lines derived from human AECs. They grow as epithelial monolayers expressing lung progenitor markers SOX9 and SOX2, with little to no expression of mature AEC markers. Co-cultured in 3-dimensions (3D) with lung fibroblasts, the cells form NKX2-1^+^ organoids expressing mature AEC markers AQP5 and GPRC5A. Single-cell RNA sequencing of an AEC line in 2D *versus* 3D revealed increased cellular heterogeneity and induction of cytokine and lipoprotein signaling, consistent with organoid formation. Activating WNT and FGF pathways resulted in larger organoids. Our approach appears to yield lung progenitor lines that retain a genetic and structural memory of their alveolar cell lineage despite long-term expansion and whose differentiation may be modulated under various 3D conditions. These cell lines provide a valuable new system to model the distal lung *in vitro*.

## INTRODUCTION

Diseases affecting the distal lung, including lung cancer, chronic obstructive pulmonary disease (COPD), idiopathic pulmonary fibrosis (IPF), and pulmonary viral infections are a significant health burden across communities worldwide. In the United States, lung cancer is still the leading cause of cancer-related deaths (Siegel et al., 2020) and chronic lower respiratory diseases account for approximately 5.7% of all deaths (Centers for Disease Control and Prevention, 2018). With the recent outbreak of COVID-19, respiratory emergencies are at an all-time high, greatly surpassing those during the 2003 SARS-CoV outbreak and the 2009 H1N1 swine flu pandemic (Centers for Disease Control and Prevention, 2020; World Health Organization, 2020; World Health Organization, 2003; Shrestha et al., 2011; Simonsen et al., 2013; Dawood et al., 2012). Because of the susceptibility of the distal lung to severe damage and the current state of public health, there is an urgent need to generate *in vitro* models to study mechanisms underlying distal lung diseases and rapidly screen for novel therapeutics.

The respiratory epithelium is lined by a variety of epithelial cell types, each with distinct functions. Along the airways, mucous-secreting goblet cells and ciliated, club, and basal cells form a pseudostratified columnar epithelium that traps and removes inhaled foreign particles (Rackley and Stripp, 2012; Tata and Rajagopal, 2017). The bronchiolar epithelium ends in saccular respiratory units, the alveoli. Alveoli are composed of two epithelial cell types: cuboidal alveolar type 2 cells (AT2) responsible for producing and secreting surfactant, and large, delicate type 1 cells (AT1) enveloped by a network of capillaries to allow gas exchange (Tata and Rajagopal, 2017; Rock and Hogan, 2011; Rackley and Stripp, 2012). AT2 cells have been extensively investigated due to their role as stem cells in the regeneration of damaged lung epithelium (Khalil et al., 1994; Liu et al., 2011; Barkauskas et al., 2013). AT2 cells can both proliferate and differentiate into AT1 cells in response to lung injury (Evans and Bils, 1969; Adamson and Bowden, 1974; Beers and Morrisey, 2011). In contrast, the fragile AT1 cells have been less extensively studied due to the difficulty in purifying and culturing these cells. Until recently, AT1 cells were thought to be terminally differentiated (Evans and Hackney, 1972; Hackney et al., 1975). However, recent reports suggest that AT1 cells, or at least a subpopulation thereof, may have greater cellular plasticity than previously thought, having the ability to reenter the cell cycle and differentiate into AT2 cells (Borok et al., 1998; Jain et al., 2015; Wang et al., 2018; Yang et al., 2016; Little et al., 2019). Thus, AT1 cells may also play a role in alveolar regeneration, albeit a more limited one.

In the last 30 years, numerous human lung cell lines have been used to study lung disease and homeostasis *in vitro*. The most common cell models available are immortalized primary airway epithelial cells. These lines were established using viral and non-viral immortalization methods aimed at overcoming cellular senescence and crisis, events marked by growth arrest and telomeric attrition, respectively (Reddel, 2000; Campisi, 2013; Herranz and Gil, 2018; Hahn and Weinberg, 2002). Human bronchial epithelial cells were first immortalized with viral proteins Simian virus 40 Large T antigen (SV40 LgT) or human papillomavirus (HPV) E6 and E7 in conjunction with the human catalytic subunit of telomerase (hTERT). BEAS-2B and small airway (SA) cells are well-known examples of such immortalized bronchial epithelial cells (Reddel et al., 1988; Lundberg et al., 2002; Piao et al., 2005; Schiller et al., 1994; Zabner et al., 2003). Subsequent non-viral methods using hTERT combined with expression of either a mutant cyclin-dependent kinase 4 (CDK4^R24C^) that is insensitive to p16^INK4A^ and p15^INK4B^ inhibition, or knockdown of p16^INK4A^/p19^ARF^ (at the *CDKN2A* locus) gave rise to the human bronchial epithelial cell line, HBEC (Ramirez et al., 2004), and the small airway epithelial cell line, SAEC (Sasai et al., 2011; Smith et al., 2016).

In contrast to the wide selection of airway epithelial cell lines showing phenotypic and morphologic similarities to primary cells, few human alveolar epithelial cell (AEC) lines have been available. Kemp *et al*. and Kuehn *et al*. immortalized purified AT2 cells from human lung tissue using SV40 LgT plus hTERT (Kemp et al., 2008) or a cocktail of proprietary ‘immortalizing’ genes (Kuehn et al., 2016). These cells quickly lost their AT2 phenotype upon 2D culturing and resembled AT1 cells in morphology and by limited marker staining of caveolin-1 (CAV1). Bove *et al*. (2014) also attempted to subculture purified AT2 cells using media containing the small molecule ROCK inhibitor, Y-27632 along with feeder cells, but reported loss of AT2 signatures after two passages and adoption of an “AT1-like” phenotype marked by expression of aquaporin 5 (AQP5) and podoplanin (PDPN). Continued growth of these cells beyond 2 months was not reported. As an alternative approach, other laboratories have derived AECs by directed differentiation of induced pluripotent stem cells (iPSC). These iPSC-derived AECs formed at varying efficiencies and with different AT2:AT1 compositions depending on the alveolarization media used (Gotoh et al., 2014; Dye et al., 2015; Jacob et al., 2017; McCauley et al., 2017; Yamamoto et al., 2017; Kanagaki et al., 2020). When suspended in Matrigel, these cells generated structures similar to purified human AT2 cells under organotypic culture (Barkauskas et al., 2013; Zacharias et al., 2018). This morphologic property has allowed for modeling of numerous alveolar pathologies (Hiemstra et al., 2018; Heo et al., 2019; Shafa et al., 2018; Korogi et al., 2019; Leibel et al., 2019; Porotto et al., 2019; Schruf et al., 2020). However, the time and experience required to differentiate iPSCs properly towards a distal lung lineage are not trivial. It takes a minimum of 50 days to generate iPSC-derived AECs (Gotoh et al., 2014; Dye et al., 2015; Jacob et al., 2017). Furthermore, improper maintenance of human fibroblasts and iPSCs can reduce conversion efficiency (Meng et al., 2012; Volpato and Webber, 2020). A simple method for deriving long-term proliferating human AECs that are rapidly expandable under standard culturing conditions and retain the ability to form functional 3D alveolar structures would be of great value to the scientific community.

Here, we report the establishment and characterization of a collection of immortalized cell lines derived from isolated adult human AT2 cells. Under 2D culture conditions, the cells grow as an epithelial monolayer and exhibit lung progenitor-like expression patterns. In 3D organotypic culture, they form diverse organoid structures and express mature AEC markers, AQP5 and GPRC5A. Cell lines derived from the alveolar epithelium are essential for the study of diseases arising from lung alveoli as they are best suited to recapitulate the pathogenesis of disease. Our novel AEC lines may therefore serve as a powerful *in vitro* model for studying genetic and environmental mechanisms underlying distal lung diseases as well as for investigating the regulation of alveolar epithelial cell homeostasis.

## RESULTS

### Direct transduction of isolated primary adult human AT2 cells does not result in immortalized epithelial cells

Primary human AECs do not proliferate in culture (Isakson et al., 2002; Mao et al., 2015), making them challenging to manipulate and limiting their suitability for *in vitro* modeling. AT1 cells are too fragile to purify and culture, therefore, we focused on the immortalization of primary human AT2 cells.

We initially attempted direct transduction of both freshly isolated and previously cryopreserved human AT2 cells with lentiviruses carrying CDK4^R24C^ and hTERT (**Figure 1 – Supplement 1** and **Figure 1 – Supplement 2A-C**) based on the reported success in human bronchial epithelial cells (Ramirez et al., 2004). This approach resulted in early proliferation of cells, followed by either growth arrest or adoption of a fibroblast-like morphology at later passages (**Figure 1 – Supplement 2A-C**). Re-transduction of seemingly growth arrested cells with SV40 LgT resulted in fibroblast-like cells (**Figure 1-Supplement 2A, last panel**). Failure to derive proliferating AECs by direct transduction motivated us to refine our immortalization strategy. In culture, AT2 cells spontaneously transdifferentiate into AT1-like cells over the course of 6 days (Cheek et al., 1989; Dobbs, 1990; Foster et al., 2007; Fuchs et al., 2003; Marconett et al., 2013). These AT1-like cells have attenuated cell bodies and a high overall surface area, similar to AT1 cells *in vivo*. We considered that our failure to immortalize primary AECs by direct transduction with lentiviruses carrying immortalizing genes might be because cells undergoing terminal transdifferentiation may not be molecularly susceptible to respond to mutant CDK4 and hTERT. In the literature, we found that successful immortalization seemed to be possible for purified primary human cells exhibiting at least some proliferative capacity (Herbert et al., 2002; O’Hare et al., 2001). Indeed, primary human bronchial epithelial cells were first subcultured before immortalization with retroviruses (Ramirez et al., 2004). We therefore hypothesized that successful immortalization of primary AECs would require first stimulating the cells to proliferate in culture, then transducing the dividing cells with lentiviruses carrying immortalizing genes (**Figure 1 – Supplement 2D**).

### Optimization of culture media conditions to propagate primary adult human AECs

To determine optimal cell expansion conditions, we performed a small-scale screen of media containing growth factors and small molecules reported to promote survival and proliferation of primary epithelial cells (**Figure 1 – Source Data 1)**. Using previously cryopreserved isolated human AT2 cells from Lung-FT (**Table 1**), we cultured approximately 6250 cells per well of a 96-well multiplate in each media condition. In the first two days after plating, most cells did not attach, and growth media contained noticeable levels of cellular debris. Wells containing ROCK inhibitor (Y-27632), keratinocyte growth factor (KGF or FGF7), 6’-bromoindirubin-3’-oxime (BIO), and 20% fetal bovine serum (FBS) showed several cells attached, but not yet flattened (data not shown). Eventually, only medium containing 10 μM Y-27632 supported cell survival and proliferation. By visual inspection, only 10-20% of the well was populated by cells. About six weeks post-plating, we passaged cells using Accutase, a gentler detachment solution compared to 0.05% trypsin-EDTA (Bajpai et al., 2008). From this passage on, cells grown in medium containing Y-27632 maintained their proliferative capacity even after detachment and re-plating. These cells (referred to as “AEC-FT-ROCKinh”) grew as a monolayer and were contact-inhibited (**Figure 1A, a**).

**Figure 1.**
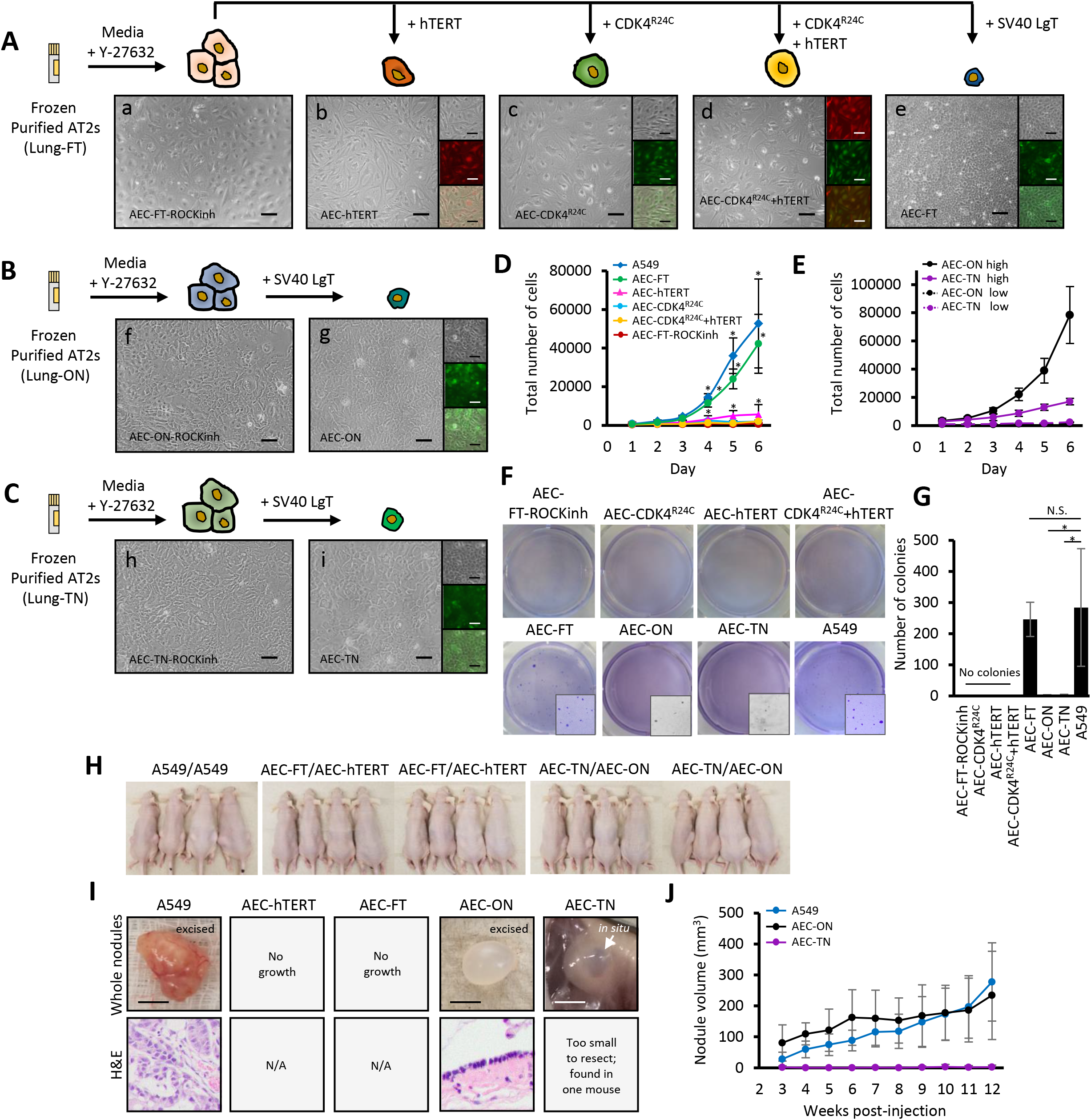
Derivation and characterization of novel alveolar epithelial cell lines. **A)** Alveolar epithelial cell line derivation scheme. Previously frozen purified human AT2 cells from Lung-FT (Table 1) were first cultured in media containing ROCK inhibitor, Y-27632, to promote cell proliferation. Proliferative cells (a) were then transduced with LeGO iT (tdTomato) and LeGO iG (eGFP) lentiviruses carrying hTERT (b), CDK4^R24C^ (c), CDK4^R24C^+hTERT (d), and SV40 LgT (e). Single and merged fluorescence images are shown as small panels on the right. Scale bar, 100 μm. Additional SV40 LgT transduced cell lines were established from AT2 cells purified from **B)** Lung-ON and **C)** Lung-TN, derived in the same manner as the schematic in (A), first by expanding in ROCK inhibitor media (f and h), then transducing with SV40 LgT (g and i). **D)** Cell proliferation assay to determine growth kinetics. One thousand cells were seeded per well in technical quadruplets and counted every day for 6 days. Plotted values are mean total number of cells ± standard deviation, n = 3 independent experiments. Differences in cell growth relative to AEC-FT-ROCKinh cells on days 4, 5, and 6 were calculated by the independent two-sample t test. Statistical significance was attained for A549 (D4: p=0.0005, D5: p=0.004, D6: p=0.02), AEC-FT (D4: p=0.002, D5: p=0.003, D6 p=0.01), and AEC-CDK4^R24C^ (D4: p=0.007, D5: p=0.005, D6: p=0.03) cells. *asterisk, p <0.05. **E)** Cell proliferation assays for AEC-LgT replicate lines, AEC-ON and AEC-TN, under standard (1000 cells/well) and high density (5000 cells/well) conditions. **F)** Representative whole well images of anchorage-independent growth assays. Five thousand cells were plated in soft agar with ROCK inhibitor growth media. A549 human lung adenocarcinoma cancer cell line, positive control. Colonies were stained with crystal violet and counted using Image J/Fiji after 1 month. Inset images are 2.5X magnified to show colonies. **G)** Colony quantification based on 6 technical replicates from at least 3 independent experiments. Plotted values are centered on mean ± standard deviation. N.S., not significant, *asterisk, p<0.05 by nonparametric Wilcoxon test. **H)** Tumorigenicity of AEC-FT, AEC-ON, and AEC-TN cell lines was assessed by subcutaneous injection into 6-week old nude (NU/J) mice for 3 months. A549 cells were used as positive control; AEC-hTERT line was used as negative control. Equal numbers of male and female mice were used in each group. Sample labels indicate which cell line was injected into the left and right flanks (Left/Right). **I)** Photos of excised nodules (scale bar, 5 mm) and corresponding H&E stainings. **J)** Graph of nodule growth starting at 3 weeks post-injection.

**Table 1.** De-identified lung donor information.

| Lung Sample | Age (years) | Sex | Race/Ethnicity | Disease Status | Cause of Death | Tobacco Use |
| --- | --- | --- | --- | --- | --- | --- |
| Lung-FT | 25 | M | Caucasian | Normal | Head Trauma | Non-smoker* |
| Lung-ON | 66 | F | Caucasian | Normal | ICH-Stroke | Non-smoker |
| Lung-TN | 62 | M | Caucasian | Normal | Stroke | Former smoker** |

ICH = Intracerebral hemorrhage
\* Patient reported as non-smoker, but gross lung displayed black speckles/particulate matter suggesting possible inhalant exposure
\*\* Patient smoked 1 pack per day for 18 years, then quit in 1991

### Derivation of a collection of proliferative human AEC lines

Although growth in medium containing Y-27632 greatly expands the lifespan of primary human cells (Liu et al, 2012; Horani et al., 2013; Chapman et al., 2014), this proliferative state is conditional and is not sufficient for immortalization (Martinovich et al., 2017; Suprynowicz et al., 2012; Bove et al., 2014). Viral oncogenes, such as HPV 15/16, EBV, and SV40 LgT have been shown previously to immortalize human cells even in the absence of hTERT (Band et al., 1991; Liu et al., 2006; Reddel et al., 1988; Neufeld et al., 1987; Christian et al., 1987). To maintain the proliferative potential of the derived AEC-FT-ROCKinh cells from Lung-FT, we transduced these cells with different combinations of lentiviruses: hTERT alone, CDK4^R24C^ alone, CDK4^R24C^+hTERT, and SV40 LgT alone (**Figure 1A**). We passaged transduced cells with Accutase and maintained them in media containing ROCK inhibitor. Epithelial-like cells resulted following each transduction (**Figure 1A, a-e**). Cells appeared proliferative, but at full confluence, formed a contact inhibited, cobblestone monolayer, suggesting maintenance of an epithelial identity. We noted that cells transduced with SV40 LgT were compact compared to cells transduced with hTERT alone, CDK4^R24C^ alone, or CDK4^R24C^+hTERT. We also observed that the SV40 LgT-transduced cells (henceforth referred to as AEC-FT) were much more rapidly expandable in culture compared to the other lines. We were easily able to maintain these cells with routine passaging for at least one year. We therefore generated biological replicates of the AEC-FT cell line from two additional human donor lungs (**Table 1**). The two additional SV40 LgT-transduced (AEC-LgT) cell lines, termed AEC-ON (from Lung-ON) and AEC-TN (from Lung-TN), exhibited epithelial-like morphology and were contact-inhibited upon confluence (**Figure 1B, C**).

### Characterization of the transformation state of human AEC lines

Since these AEC lines were generated using chemical and genetic modifications, it was possible that the cells had acquired additional changes precluding their use as “normal” cell models. To determine whether the newly derived cells might have acquired any characteristics of oncogenic transformation, we performed proliferation and soft agar assays. As a reference for full cellular transformation, we used the lung adenocarcinoma cell line, A549 (Giard et al., 1973). We assessed the rate of proliferation by seeding cells into a multiwell plate and counted the total number of cells every day for six days. On days 4, 5, and 6, we noted statistically significant differences in the number of A549, AEC-FT, and AEC-CDK4^R24C^ cells compared to AEC-FT-ROCKinh cells (**Figure 1D**). AEC-FT cells proliferated faster than AEC-FT-ROCKinh, AEC-hTERT, AEC-CDK4^R24C^, and AEC-CDK4^R24C^+hTERT cells, reaching exponential growth around day 4 (**Figure 1D**). To determine whether the extended lag phase of the slow-growing cells was due to initial seeding density or truly reflective of the cell’s replicative capacity, we performed the same proliferation assay with a higher initial seed number of 5000 cells. Under these conditions, AEC-FT-ROCKinh, AEC-hTERT, AEC-CDK4^R24C^, and AEC-CDK4^R24C^+hTERT cells still did not achieve exponential growth; therefore population doubling times (PDTs) were not calculated (**Figure 1 – Supplement 3A, B**). AEC-ON and AEC-TN cell lines exhibited a similar extended lag phase to AEC-FT-ROCKinh, AEC-CDK4^R24C^, AEC-hTERT, and AEC-CDK4^R24C^+hTERT cells and did not reach exponential growth (**Figure 1E**). However, when plated at a higher density, cells proliferated readily, reaching exponential growth 4 days after plating, with PDT of 1.1 ± 0.05 days for AEC-ON and 2.1 ± 0.4 days for AEC-TN (**Figure 1 – Supplement 3B**).

To determine whether the cells exhibited anchorage-independent growth, a common feature of transformed cells, we performed soft agar assays on all AEC lines, suspending 5000 cells in agarose melted in growth media containing ROCK inhibitor. We assessed colony formation after one month by crystal violet staining using A549 cells as a positive control (**Figure 1F**). No colonies were detected for AEC-FT-ROCKinh, AEC-hTERT, AEC-CDK4^R24C^, or AEC-CDK4^R24C^+hTERT cells. In contrast, AEC-FT cells consistently generated soft agar colonies (246 ± 55 colonies per well), comparable to A549 cells (284 ± 186 colonies; the difference was not significant by Wilcoxon nonparametric test, p-value = 0.78) (**Figure 1G**). AEC-ON and AEC-TN cells did not form appreciable colonies; rare small colonies were visible under 10X brightfield magnification (**Figure 1F, inset**). Quantification of colonies yielded 2 ± 1 colonies per well for AEC-ON cells and 3 ± 2 colonies for AEC-TN, which was statistically significantly different from A549 cells (p = 1.4 × 10^−7^ and p = 1.5 × 10^−7^, respectively, by nonparametric Wilcoxon test) (**Figure 1G**).

### AEC-LgT cell lines exhibit different growth potentials in nude mice

We noted that the BEAS-2B cell line, derived from bronchial epithelial cells immortalized using SV40 LgT, also form colonies in soft agar, but do not form tumors in nude mice. We therefore subcutaneously injected the three AEC-LgT lines (AEC-FT, AEC-ON, AEC-TN) into nude mice to test cell tumorigenicity. We used AEC-hTERT cells as a negative control because they were moderately proliferative out of the four slow growing cell lines (see Figure 1D), but did not form colonies in soft agar. A549 cells, which are known to form colonies in soft agar (Jiang et al., 2001) and tumors in mouse xenografts (Kang et al., 2006), were used as a positive control. We injected 1×10^6^ cells of each cell line suspended in Matrigel into the flanks of nude mice and monitored the size of growths beginning 3 weeks post-injection. After 3 months, mice transplanted with A549 cells and AEC-ON cells displayed prominent nodules (**Figure 1H**). Excised nodules were evaluated by an expert pathologist blind to sample identities. A549-derived nodules were solid tumors of adenocarcinoma histology, whereas AEC-ON-derived nodules were non-malignant, fluid-filled cysts lined by a simple epithelium with areas of pseudostratified cells (**Figure 1I**). In one case (1 out of 8 flanks), AEC-TN cells formed a nodule less than 20 mm^3^ that was too small to resect. This nodule did not significantly change in volume during the 3-month experimental period (**Figure 1J**). Neither AEC-FT nor negative control AEC-hTERT cells formed nodules. Taken together, although SV40 LgT transduction conferred cell immortality, it did not appear to induce malignant transformation.

### AEC lines are transcriptomically distinct from primary human AECs and LUAD cell lines

To determine how similar the transcriptomic identities of our novel alveolar epithelial cell lines are to their parental purified primary AECs, we compared their transcriptomes to those of primary AECs by paired-end RNA-sequencing. We also compared our AEC lines to lung fibroblasts to test for epithelial-to-mesenchymal transition gene signatures, as well as to LUAD cancer cell lines to determine if our novel cells bore transcriptional hallmarks of oncogenic transformation. We isolated total RNA from our novel AEC lines and the human lung fibroblast cell line HLF-133 cultured at subconfluence. For the primary AEC sample group, total RNA was harvested from purified human AT2 cells and AT2 cells transdifferentiated on filters for 2 (D2), 4 (D4), and 6 (D6) days into AT1-like cells (Danto et al., 1995) from three donor lungs designated “AEC-f,” “AEC-m,” and “AEC-a” (**Figure 2 – Source Data 1**). We downloaded additional RNA-seq data for lung fibroblasts and LUAD cancer cell lines from publicly available databases ENCODE (www.encodeproject.org) and DBTSS (www.dbtss.hgc.jp), analyzing a total of 42 samples (**Figure 2 – Source Data 1**).

To examine relationships between samples based on their gene expression patterns, we performed Principal Component Analysis (PCA) and sample-sample distance matrix analyses on the top 500 most variable genes across the data set. Our analyses showed six clusters: lung fibroblasts, the original AEC lines from Lung-FT (5) together, the biological replicate AEC-LgT lines from Lung-ON and -TN, fetal lung tissue, LUAD cancer lines, and primary AECs (**Figure 2A**). Human fetal lung tissue clustered away from all samples. Although the biological replicate lines AEC-ON and AEC-TN were generated by transduction of SV40 LgT in a similar fashion to AEC-FT cells, these two lines clustered separately from the original collection of ROCKinh-derived cells from Lung-FT. Thirty-nine percent of the variation in the data captured by principal component 1 (PC1) divided the entire sample set into clusters containing long-term cultured cells (LUAD, fibroblasts, AEC lines) and the other cells (primary AECs and fetal lung tissue), underlining the possible effects of growing cells in culture. Groups identified in the sample distance matrix were generally in agreement with the PCA analysis; however, lung fibroblasts were found to be grouped in a distinct cluster from AEC lines based on dissimilarity calculations represented by a dendrogram tree (**Figure 2B**).

**Figure 2.**
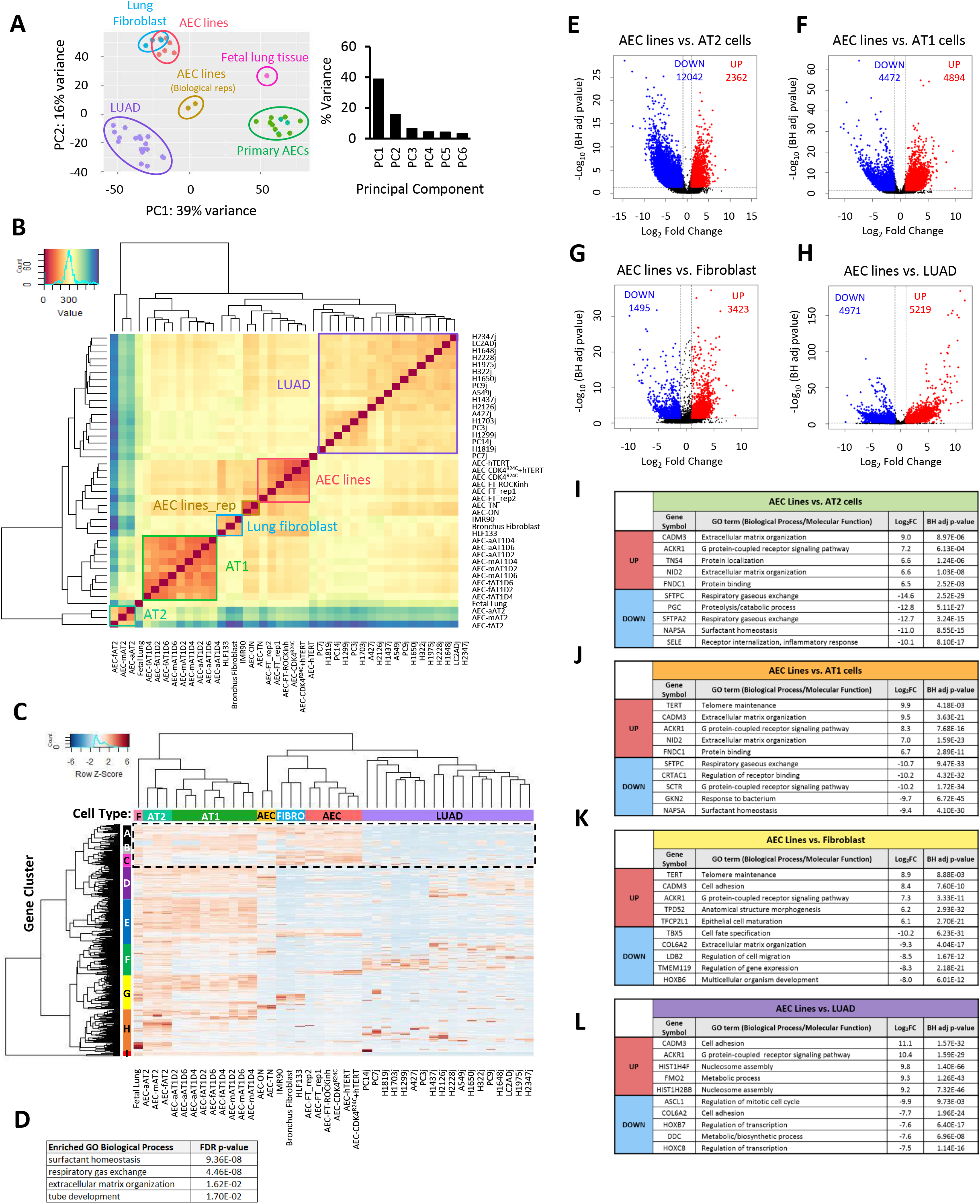
Exploratory and differential gene expression analyses of transcriptomic profiles of AEC lines, lung fibroblasts, primary AECs, and LUAD cell lines reveal widespread gene expression differences. **A)** Left: PCA plot showing 6 groupings based on the top 500 most variable genes using DESeq2 pipeline. Each sample is represented by a dot and colored according to sample type. Right: Bar chart of percent variance across principal components (PCs) showing the majority of the variance in gene expression within the dataset is described by the first two PCs. **B)** Sample-sample distance matrix showing 6 sample groups based on similarity calculations using Euclidean distance metric. Dark red diagonal indicates complete similarity between identical samples. Blue hues indicate low similarity. Primary human AT2 cells are labeled as “AEC-…AT2,” with “f,” “m,” and “a” indicating three separate donor lungs. Primary human AT1-like cells *in vitro* differentiated from AT2 cells are labeled as “—D2,” “—D4, “—D6,” with numbers representing days transdifferentiated in culture. **C)** Unsupervised hierarchical clustering on the the top 500 most variable genes. Cell type categories are indicated by the horizontal colored bars: F, fetal lung tissue; AT2, primary human AT2 cells; AT1, human AT1-like cells; AEC (yellow), AEC-ON and AEC-TN replicate cell lines; Fibro, human lung fibroblasts; AEC (pink), AEC line collection derived from Lung-FT; LUAD, human lung adenocarcinoma cancer cell lines. Gene clusters identified from the dendrogram are shown by vertical colored bars, labeled A-I. **D)** GO terms corresponding to Gene Clusters A, B, and C (dotted line) which differentiate primary AECs, lung fibroblasts, and AEC lines from LUAD cancer lines. GO terms were determined by PANTHERv14.1 Overrepresentation Test (Released 20200728) using Fisher’s Exact Test from the Gene Ontology Consortium. Statistical significance cutoff was set to FDR-corrected p-value < 0.05. Differential gene expression analyses were performed between all derived AEC lines and: **E)** AT2, **F)** AT1-like, **G)** lung fibroblasts, and **H)** LUAD cancer cell lines and shown as Volcano plots. Statistically significant differentially expressed genes relative to AEC lines are shown in **blue**(**downregulated**) and **red**(**upregulated**). Significance cut offs (dotted lines) were set at absolute value of Log_2_ Fold Change ≥ 1 (therefore |Fold Change| ≥ 2) and Benjamini-Hochberg (BH) adjusted p-value < 0.05; x-axis, Log_2_Fold Change; y-axis, −Log_10_BH adjusted p-value. **I-L)** Table of GO terms for the top 5 most significantly upregulated and 5 most significantly downregulated protein-coding genes. Listed Log_2_Fold Change and BH adjusted p-values are from differential gene expression analyses.

To determine which genes drive segregation of the different sample clusters, we performed unsupervised hierarchical clustering using the top 500 most variable genes across the data set. Consistent with PCA and sample distance matrix plots, primary AECs and fetal lung tissue were more related to each other than to the remaining groups of samples, as shown in the column dendrogram (**Figure 2C**). Nine salient gene clusters were identified within the top 500 genes (**Figure 2 – Supplement 1A**). Most of the clusters had at least one Gene Ontology (GO) term related to tissue development, morphogenesis, and cellular response to the environment (**Figure 2 – Supplement 1B**). Inspection of the differences in overall gene expression across the sample types (fetal lung tissue, primary AT2, *in vitro*-derived AT1-like, AEC lines, lung fibroblasts, and LUAD) revealed a group of genes with higher expression in non-cancer samples compared to LUAD samples (**Figure 2C, dashed box**). Genes associated with this group had GO terms for “surfactant homeostasis,” “regulation of immune system process,” “response to external biotic stimuli,” and “tube development” (**Figure 2D**). Highlighted in Figure 2D, gene clusters A, B, and C distinguished fetal lung tissue, primary AECs, lung fibroblasts, and AEC lines from LUAD cancer cell lines and were enriched in genes involved in normal lung physiology (**Figure 2 – Source Data 2**).

We next performed differential gene expression analyses between the AEC lines and each of the other groups: AT2 cells, AT1-like cells, lung fibroblasts, and LUAD cancer lines. Volcano plots for each comparison were generated to visualize statistically significant differentially expressed genes. Comparing AEC lines to primary AT2 cells, a total of 14,404 genes were differentially expressed, the largest number in the four pairwise comparisons (**Figure 2E**). Compared to AT1-like cells, 9366 total genes were differentially expressed in AEC lines (**Figure 2F**). Comparing the AEC lines and lung fibroblasts, 4918 genes were significantly differentially expressed, the least number of gene differences of the four pairwise comparisons (**Figure 2G**). Comparing the AEC lines to LUAD cancer lines, a total of 10,190 genes were significantly differentially expressed (**Figure 2H**). Notably, although the range of fold change values for these genes was similar to the fold change ranges for the other comparisons (Log_2_FoldChange axis between −15 and 15, shown in **Figure 2E-H**), the −Log_10_ BH adjusted p-values associated with the top upregulated genes were orders of magnitude more significant (**Figure 2H** y-axis compared to y-axes in **2E-G**). Examining GO terms of the top upregulated genes between the AEC lines and both AT2 and AT1-like cells, we found genes related to alterations in cell adhesion and extracellular matrix signaling, in line with the altered cell morphology and proliferative potential of the AEC lines compared to primary AECs. GO terms associated with downregulated genes in the AEC lines compared to primary AECs were related to respiratory and surfactant homeostasis (**Figure 2I-J**). When compared to lung fibroblasts, upregulated genes in the AEC lines were associated with telomere maintenance and morphogenesis. Downregulated genes were associated with cell migration and extracellular matrix organization (**Figure 2K**). GO terms for top upregulated genes in AEC lines *versus* LUAD cancer lines corresponded to cell signaling, metabolism and nucleosome remodeling, indicating significant differences between non-cancer- and cancer-derived cell lines. For the top downregulated protein-coding genes, associated GO terms were generally related to cell cycling and transcription (**Figure 2L**).

Based on differential gene expression analyses and PCA clustering, the AEC lines appeared to be more similar to lung fibroblasts than to primary AECs, suggesting a possible loss in lung epithelial cell-specificity and adoption of a more mesenchymal phenotype. To determine whether the AEC lines retained their lung-specific gene expression patterns, we subset the expression data on 75 lung-related genes manually curated from published bulk RNA-seq and single-cell RNA-seq data (Treutlein et al., 2014; Xu et al., 2016) (**Figure 2 – Source Data 3**). As shown in **Figure 2 – Supplement 2A**, compared to purified primary AECs, expression of these lung-related genes was low in the AEC lines. However, compared to lung fibroblasts, the AEC lines expressed these genes at a much higher level (**Figure 2 – Supplement 2B**). This suggests that the similarity between the AEC lines and fibroblasts calculated by PCA analysis could be driven by the fact that fibroblasts are the only other cell type that was non-cancer, of normal ploidy, and cultured on plastic.

### AEC-LgT cells express features of lung progenitor cells

The ROCK inhibitor Y-27632 is a commonly used small molecule to promote stem cell survival and proliferation (Claassen et al., 2009; Vernardis et al., 2017). The addition of ROCK inhibitor and feeder cells has also been shown to enhance culturing of primary epithelial cells from mammary, prostate, and upper airway lung tissues (Liu et al., 2012). However, in this process of facilitating cell survival, adult cells are reprogrammed to a stem-like state (Suprynowicz et al., 2012). In the mouse lung, SOX9 regulates distal lung cell fate, committing early cells to an alveolar epithelial cell lineage (Rockich et al., 2014; Chang et al., 2013), whereas SOX2 is an important regulator of proximal lung cell fate, committing early lung stem cells to the basal cell lineage (Ochieng et al., 2014; Daniely et al., 2004). In the human lung, however, recent reports described a new population of progenitors, located at budding distal epithelial tips during fetal lung development, that co-express SOX9 and SOX2 (Danopoulos et al., 2018; Nikolić et al., 2017). We therefore assessed the expression of lung progenitor markers SOX9 and SOX2 in the three AEC-LgT lines by immunofluorescence (IF) staining and found that all three co-expressed SOX9 and SOX2 proteins (**Figure 3A**). In agreement with the IF staining, RNA-seq showed the AEC lines, as a group, expressed both SOX9 and SOX2 genes more highly than primary AECs (**Figure 3 – Supplement 1**).

**Figure 3.**
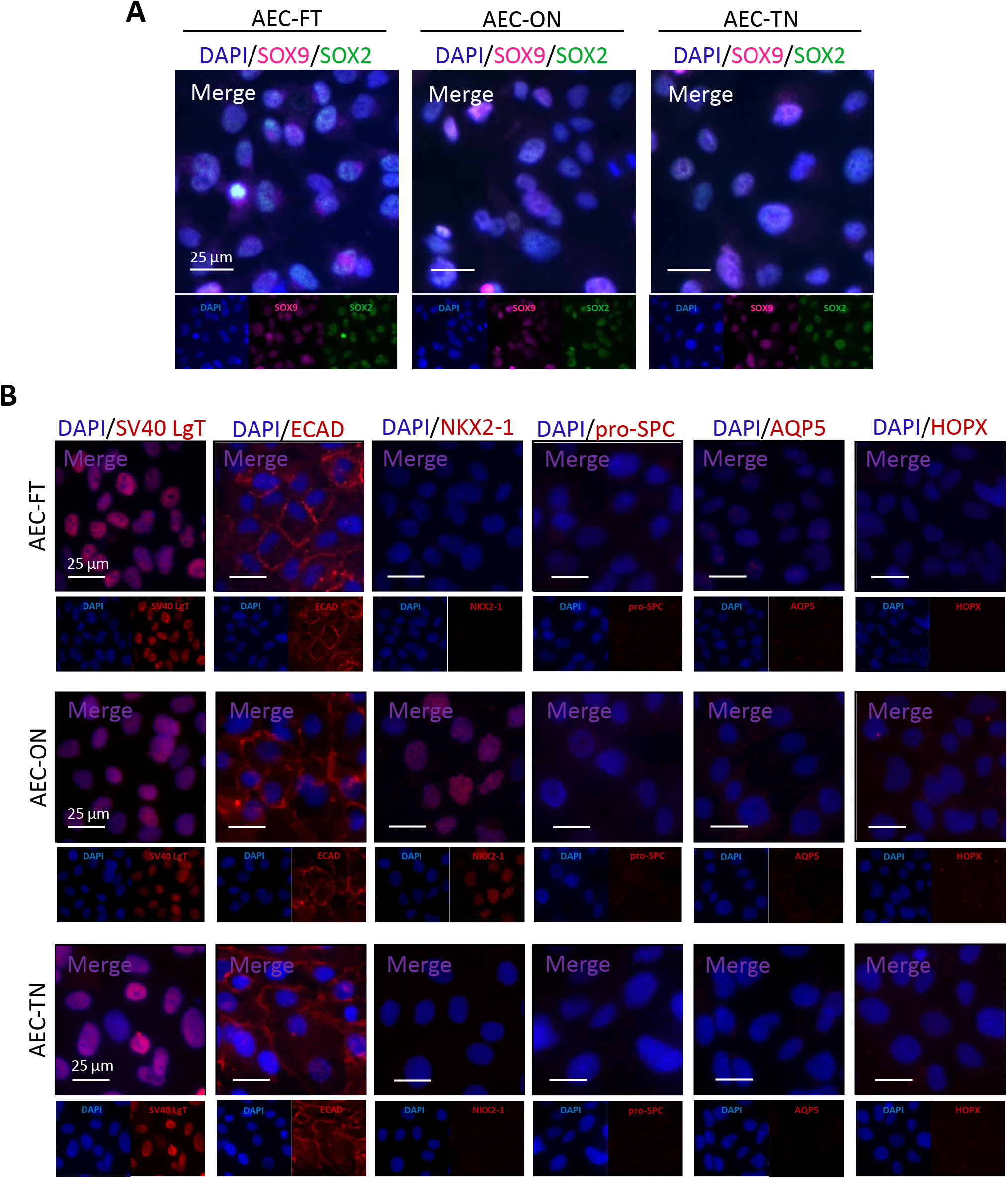
Alveolar epithelial cell lines prominently express lung progenitor markers over mature AEC markers. Merged and single channel images of representative immunofluorescence staining of AEC-LgT cells grown on standard culture dishes. **A)** Co-staining of SOX9 and SOX2 lung progenitor markers for AEC-FT, AEC-ON, and AEC-TN cell lines. **B)** Staining of SV40 LgT and epithelial marker, E-cadherin (ECAD), and mature AEC markers NKX2-1, pro-SPC, APQ5, and HOPX. Staining patterns were consistent across different passages. At least three independent experiments were performed. Scale bar, 25 μm.

To examine the expression status of mature lung markers in the three AEC-LgT cell lines, we again performed IF staining on monolayer cultures. We first determined that all AEC-LgT lines were positive for SV40 LgT and E-cadherin (ECAD), an epithelial cell-cell junction protein, indicating the cells retained their epithelial cell phenotype. All three cell lines were negative for mature AT1 cell markers AQP5 and homeodomain-only protein (HOPX), and AT2 cell marker pro-surfactant C (pro-SPC). AEC-FT and AEC-TN were negative for NKX2-1 (also known as thyroid transcription factor-1, TTF1). In contrast, AEC-ON cells expressed NKX2-1, although at variable levels across the cell population (**Figure 3B**). NKX2-1 is a central transcriptional regulator of lung endoderm specification, branching morphogenesis, and both proximal and distal lung cell differentiation during development (Herriges and Morrisey, 2014; Minoo, 2000; Yuan et al., 2000). Patients with NKX2-1 haploinsufficiency exhibit recurring pulmonary complications and respiratory distress, among other physiological dysfunctions (Hamvas et al., 2013). Notably, although NKX2-1 regulates expression of the AT2 cell marker surfactant protein C (SFTPC) (Kelly et al., 1996), we did not detect expression of SFTPC in AEC-ON cells, as marked by its precursor protein, pro-SPC. Taking our transcriptomic analyses and IF staining data together, we speculated that AEC-LgT cells in 2D culture favored a transcriptional program promoting cell proliferation and cell survival over one specifying alveolar epithelial cell lineage. We therefore investigated whether, under certain conditions, AEC-LgT cells could be induced to differentiate to a phenotype resembling human adult AECs.

### AEC-LgT cells can form lung organoids in 3D co-culture

Purified AT2 cells from mouse and human lungs have been shown to form organoids when cultured with stromal cells and suspended in Matrigel (Barkauskas et al., 2013; Jain et al., 2015; Zhou et al., 2018; Zacharias et al., 2018). This 3D culture system has also been used to assess differentiation capabilities of both AT2 progenitor cells and iPSC-derived distal lung cells (Jacob et al., 2017; McCauley et al., 2017; Yamamoto et al., 2017). To determine whether AEC-LgT cells possess the ability to form 3D structures, we mixed exponentially-growing AEC-FT, AEC-ON, and AEC-TN cells with neonatal mouse fibroblast cells (MLg) in Matrigel and co-cultured them in Transwell dishes (**Figure 4A**). All three cell lines formed organoids from a single-cell suspension (**Figure 4B**), in contrast to the dense cell clusters formed by A549 cells under similar conditions (**Figure 4 – Supplement 1**). In the absence of MLg fibroblasts, we did not observe organoid formation from any of the AEC-LgT cell lines after 1 month of culture (**Figure 4 – Supplement 2**).

**Figure 4.**
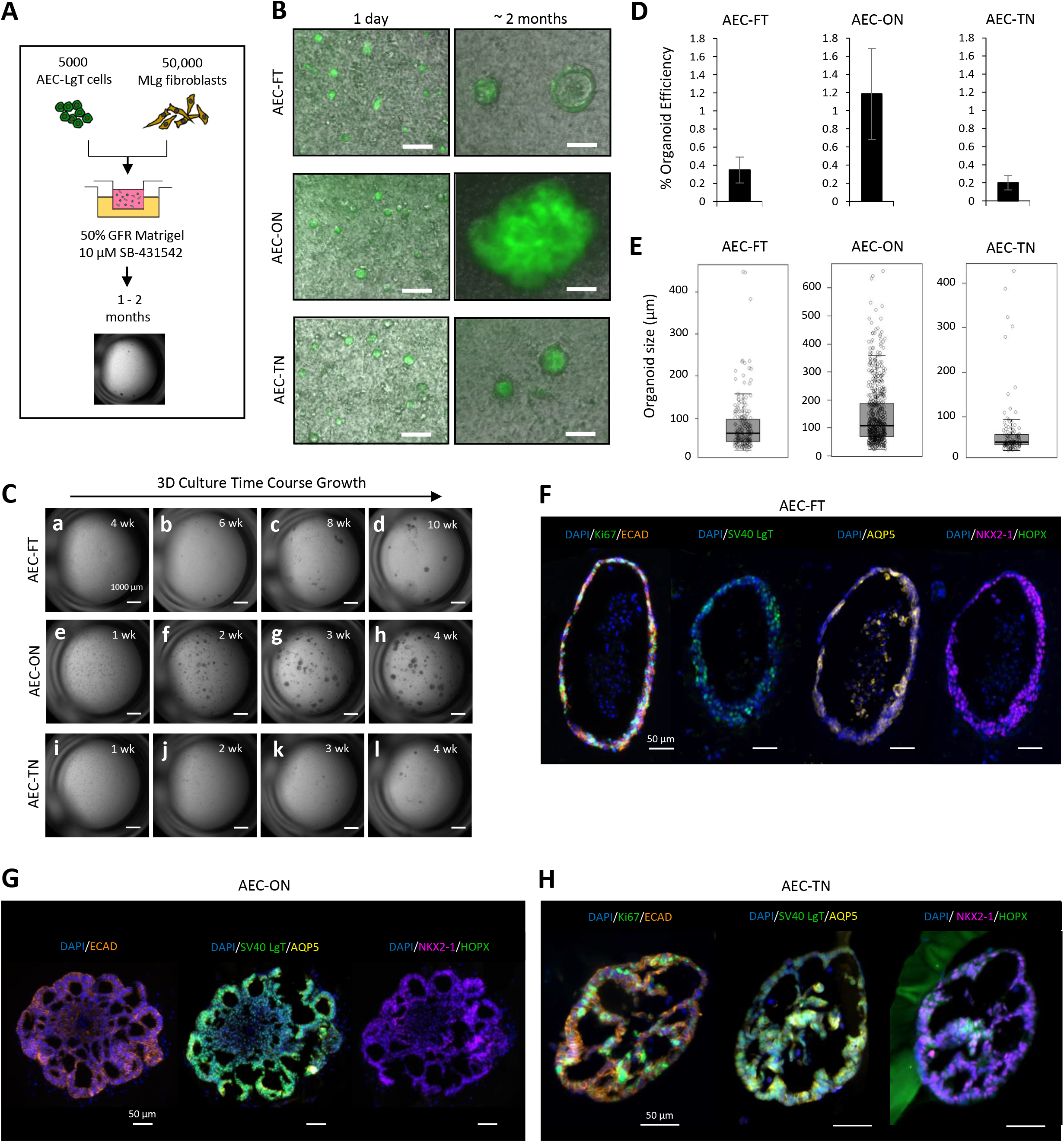
AEC-LgT cell lines form organoids in 3D co-culture and express mature lung markers. **A)** AEC-LgT cells were co-cultured with MLg mouse neonatal fibroblasts in growth factor reduced (GFR) Matrigel on Transwell inserts surrounded by media containing 10 μM SB-431542 for at least 1 month. **B)** Organoids arise from a single cell suspension of SV40 LgT^+^ epithelial cells, as determined by presence of GFP. Scale bar, 100 μm. **C)** Time course growth of AEC-FT, AEC-ON, and AEC-TN organoids. AEC-FT organoids grew much slower than either AEC-ON or AEC-TN organoids, therefore time course imaging began at 4 weeks, with subsequent imaging every 2 weeks (a-d). AEC-ON (e-h) and AEC-TN (i-l) cultures were imaged at 1 week until 4 weeks, imaging every week. Scale bar, 1000 μm. **D)** Organoid formation efficiencies and **E)** organoid size measurements for AEC-FT, AEC-ON, and AEC-TN cells at 2 months, n = 3 biological replicates, each with at least 6 inserts. Immunofluorescence staining of representative organoid sections from **F)** AEC-FT, **G)** AEC-ON, and **H)** AEC-TN cell lines after 1 month of growth. Organoids were confirmed to retain epithelial characteristics by SV40 LgT and ECAD staining. Ki67, proliferation marker; AQP5, HOPX, NKX2-1, mature AEC markers. Scale bar, 50 μm.

Mouse and human AT2 cells form organoids within several weeks of culturing (Jain et al., 2015; Barkauskas et al., 2013; Zacharias et al., 2018), whereas SOX9^+^/SOX2^+^ human distal tip progenitors form organoids in as early as 12 hours (Nikolić et al., 2017). To examine the rate of organoid formation of all three AEC-LgT cell lines, we carried out a time-course experiment. AEC-FT cells were noticeably slower in forming organoids than either AEC-TN or AEC-ON cells (**Figure 4C**), requiring at least 5 weeks to form detectable spheres at 2.5X magnification on a stereomicroscope (**Figure 4C, b**), compared to 3-4 weeks for AEC-TN (**Figure 4C, k-l**), and < 2 weeks for AEC-ON cells (**Figure 4C, f**). We noted that the rates of organoid formation for the three AEC-LgT cell lines did not coincide with their rates of proliferation in 2D culture, suggesting this structural change was not merely determined by cell division.

To quantitatively characterize organoid growth of the AEC-LgT lines, we calculated organoid formation efficiency and size after two months of culture. Organoid formation efficiency was defined as the total number of organoids divided by the initial cell seeding number (5000 epithelial cells). The mean organoid formation efficiency for AEC-FT was 0.35 ± 0.1%, for AEC-ON, 1.2 ± 0.5%, and for AEC-TN cells, 0.2 ± 0.08% (**Figure 4D**). Organoid size (diameter) was measured as the largest distance between two points on the organoid membrane. Only organoids greater than 20 μm diameter were considered for these measurements. The range of sphere sizes for AEC-FT was 25 – 445 μm (median 64 μm), for AEC-ON, 24 – 661 μm (median 108 μm), and for AEC-TN, 25 – 427 μm (median 44 μm) (**Figure 4E**). **Figure 4 – Source Data 1** summarizes the results.

Organoid shapes were variable across cultures for each AEC-LgT line; however general growth patterns were observed. Under brightfield microscopy, AEC-FT cells formed organoids of predominantly round morphology with a single lumen. Occasionally, across different 3D cultures of these cells, organoids containing multiple lumens were observed (**Figure 4 – Supplement 3A**). AEC-TN cells also formed rounded, single lumen organoids more commonly than multi-lumen organoids. However, the multi-lumen organoids, although rare, tended to appear more complex in structure than those from AEC-FT cells (**Figure 4 – Supplement 3C**). In contrast, AEC-ON organoids were more heterogeneous in morphology. A subpopulation of them were large and floret-like, exhibiting a lobulated structure (**Figure 4 – Supplement 3B**).

### AEC-LgT organoids robustly express alveolar epithelial markers

We found that AEC-LgT cells formed organoids in 3D culture, an ability well-documented in primary mouse and human AT2 cells grown under similar conditions. However, AT2 cell-derived organoids tend to be relatively dense with a small central lumen (Barkauskas et al., 2013; Zacharias et al., 2018; Zhou et al., 2018). AEC-LgT cell organoids, in contrast, were more reminiscent of organoids formed by multipotent distal tip progenitors (Nikolić et al., 2017), having marked spherical to lobulated morphologies and large lumens. To determine whether this morphological behavior was accompanied by changes in lung-specific marker expression, we performed IF staining for mature alveolar markers on organoid sections. Organoids from AEC-ON and AEC-TN cells were fixed after approximately one month of growth; organoids from AEC-FT cells were fixed after two months of 3D growth due to their slower rate of organoid formation.

By IF staining, organoids were composed of SV40 LgT^+^/ ECAD^+^ epithelial cells. All organoids contained Ki67^+^ proliferative cells that were distributed throughout the structure (**Figure 4F-H**). Upon probing for mature alveolar markers (NKX2-1, pro-SPC, AQP5, HOPX), we found that all organoids robustly expressed AQP5 and NKX2-1, whereas HOPX expression was variable (**Figure 4F-H** and **Figure 4 – Supplement 4A-C**), and pro-SPC was negative (**Figure 4 – Supplement 4D**). The commonly used AT2 cell-specific marker for sorting, HTII280, was also negative across all lines (**Figure 4 – Supplement 5A-B**). We also probed for the newly identified AT1 marker, G-protein coupled receptor family C, group 5A (GPRC5A) (Horie et al., 2020), finding that all three AEC-LgT organoids were positive, specifically along the apical lining of the lumens (**Figure 4 – Supplement 5B**), in contrast to the ubiquitous GPRC5A expression observed in all three monolayer cultures (**Figure 4 – Supplement 5A**). In AEC-FT (**Figure 4F**) and AEC-TN organoids (**Figure 4H**), 3D co-culture conditions appeared to reactivate AQP5 and NKX2-1 expression from their silenced state in 2D culture. In contrast, AEC-ON cells maintained NKX2-1 expression in both 2D and 3D cultures, and expressed AQP5 only in 3D culture (**Figure 4G**).

### Single-cell transcriptomic analyses of AEC-ON organoids reveal increased cellular heterogeneity in response to organotypic culture

As shown in **Figure 4G** and **Figure 4 – Supplement 4B**, AEC-ON organoids exhibited the most dramatic morphologies and marked expression of the AT1-enriched gene *AQP5*, whereas the AT2 cell marker *SFTPC* was not expressed, despite the presence of its upstream regulator *NKX2-1*. Currently, purified primary human AT2 cells have been shown to form organoids; as of yet, the same ability has not been reported for human AT1 cells. Because AEC-ON cells may possess expression profiles of AT2 and AT1 cells that are too nuanced to detect by IF staining, we investigated the transcriptomes of these cells grown in 2D and in 3D in greater detail by single-cell RNA-sequencing (scRNA-seq). AEC-ON cells grown on standard tissue culture plastic in ROCKinh media constituted the “2D” sample and one month-old AEC-ON organoids constituted the “3D” sample. AEC-ON organoids were gently detached and purified from the surrounding Matrigel by sequential rounds of Dispase protease treatment and centrifugation. Surrounding single cells that did not form organoids were excluded from the collection by modulating Dispase treatment times followed by low-speed centrifugation and visual inspection of the sample suspension. Dissociated and FACS-sorted GFP^+^ single cells derived from AEC-ON 2D and 3D organoids were then processed using the 10x Genomics Chromium platform (**Figure 5A**). Uniform Manifold Approximation and Projection (UMAP) dimensional reduction was performed on 10,965 total cells. AEC-ON cells grown under standard 2D conditions clustered separately from cells comprising lung organoids (**Figure 5B, left**). Cluster analyses revealed 7 distinct groups: one cluster (Cluster 0) encompassed all 2D cells and the remaining clusters (Cluster 1-6) were identified in the 3D sample, indicating increased cellular heterogeneity among cells comprising AEC-ON organoids *versus* cells in the monolayer population (**Figure 5B, right**). Cell numbers per cluster are shown in **Figure 5C**.

**Figure 5.**
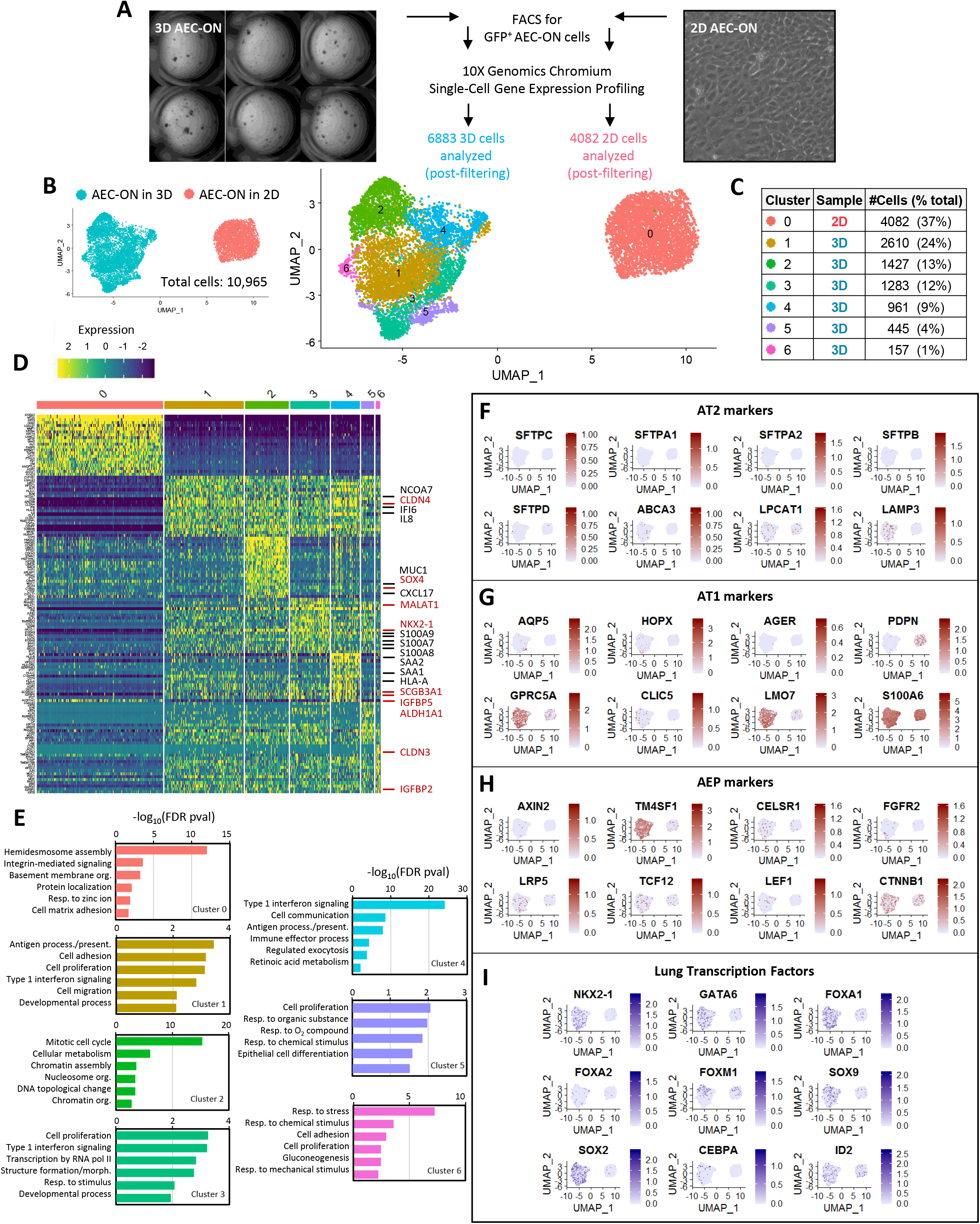
Single-cell transcriptomic analyses of AEC-ON cells reveal increased cellular heterogeneity in response to organotypic culture. **A)** Brief workflow of scRNA-sequencing experiment showing whole-well images of AEC-ON organoids and brightfield image of AEC-ON 2D cells, before dissociation and FACS isolation. Following 10x Genomics Chromium Single Cell (v3) barcoding, library prep, and sequencing, 7144 3D cells and 4320 2D cells (11,464 total cells) were pre-processed and quality controlled. After filtering out cells with low reads and a mitochondrial content of greater than 18%, a total of 10,965 cells were analyzed (6883 3D cells; 4082 2D cells). **B)** Uniform Manifold Approximation and Projection (UMAP) representation of 10,965 total cells showing grouping by sample type (left) and called clusters (right). **C)** Cell clustering proportions showing all 2D cells grouped into Cluster 0. **D)** Heat map of the top 20 gene markers enriched in each cluster compared to all cells, ordered by average expression (Log_2_FoldChange). Genes related to immune response and cytokine signaling are highlighted in **black**. Genes related to development/morphogenesis are highlighted in **red**. **E)** GO terms for the top 50 (top 75 genes, Cluster 5 only) most highly expressed genes in each cluster, as determined by PANTHERv14.1 Overrepresentation Test (released 20200728) using Fisher’s Exact Test with FDR correction for multiple testing. Only FDR values p < 0.05 were considered statistically significant and are shown in the graphs. Abbreviations: org., organization; resp., response; process., processing; present., presentation; morph., morphogenesis. **F-I)** Feature plots of selected AT2 and AT1 cell markers, alveolar epithelial progenitor (AEP) genes, and lung-related transcription factors.

To determine the set of marker genes distinguishing each called cluster, we performed differential gene expression analyses. **Figure 5D** shows a heat map of the top 20 genes in each cluster ranked by average Log_2_Fold Change over all cells. Associated GO terms for the top 50 genes ordered by p-value are shown in **Figure 5E**. For Cluster 5, GO analysis was performed on the top 75 genes, since the top 50 genes yielded no statistically significant results. Cluster 0 containing all cells from the 2D sample showed marked enrichment for genes associated with cell adhesion processes, consistent with cell growth as a monolayer (**Figure 5E**). Cluster 2 was enriched with cell cycling genes, representing actively proliferating cells within AEC-ON organoids. The remaining clusters (1, 3-6) were enriched in genes associated with response to external stimuli such as immune response, inflammation, and type 1 interferon signaling, suggesting that cells interacted with the surrounding microenvironment by secreting cytokines and lipoproteins. Clusters 1 and 3-5 were also significantly enriched for genes involved in development, morphogenesis, and epithelial cell differentiation, consistent with structural changes required for organoid formation (**Figure 5E**).

Feature maps of marker genes representative of AT2 cells, AT1 cells, and alveolar epithelial progenitor (AEP) cells (Zacharias et al., 2018) revealed a higher proportion of positive cells in 3D organoids than in 2D cultured cells (**Figure 5F-H**). Examining the AT2 cell panels, we observed numerous cells in the 3D sample expressing *ABCA3* and *LAMP3*, encoding markers of specialized organelles in AT2 cells called lamellar bodies, where surfactant is produced and stored. Consistent with expression of these genes, we detected several cells expressing surfactant proteins A2 (*SFTPA2*), B (*SFTPB*), and D (*SFTPD*) (**Figure 5F**). We did not observe a significant number of cells expressing *SFTPC*, the most lung and AT2 cell-specific surfactant produced by lamellar bodies. Examining the plots of AT1-enriched markers, we found that cells comprising the organoids highly expressed transcripts of the actin-binding protein LMO7 and, to a lesser extent, the chloride channel protein CLIC5, but expressed *AQP5* at a lower level than expected from our IF staining (**Figure 5G**), possibly due to insufficient transcript capture rate. Thus, the results we observe may be an underestimate of actual expression levels and cell distribution. Overall, the mixed population of organoid cells expressing both AT2 and AT1 genes suggests the presence of immature AT2-like cells or an AT2-AT1 intermediate cell type (Liebler et al., 2015).

AEPs were previously found to highly express the surface marker TM4SF1 and the cytoplasmic protein AXIN2, and to be WNT- and FGF-responsive (Zacharias et al., 2018). Compared to cells grown in 2D, we found a number of AEC-ON cells in 3D expressing *AXIN2* and a much greater proportion expressing *TM4SF1*. In addition, AEC-ON cells expressed genes of the WNT (*TCF12*, *LEF1*, and *CTNNB1*) and FGF pathways (*FGFR2*) (**Figure 5H**). We examined lung-related transcription factor expression in AEC-ON cells in 2D *versus* 3D, finding a higher number of cells expressing *NKX2-1*, *GATA6*, *FOXA1*, *FOXA2*, and *CEBPA* in 3D than in 2D. *SOX2* and *SOX9* were also highly expressed in 3D samples (**Figure 5I**). Since AEC-ON organoids contained AEP-like cells expressing genes related to WNT and FGF pathways, we investigated whether activation of these signals can modulate organoid growth characteristics.

### Activation of WNT or FGF signaling has cell line-specific effects on organoid growth

WNT and FGF signaling pathways are important for patterning and growth of the developing lung bud, as well as maintaining adult lung homeostasis. During lung development, WNT signaling is crucial for alveologenesis and maturation through regulation of AT2 cell self-renewal (Nabhan et al., 2018; Frank et al., 2016). FGF7 (also known as keratinocyte growth factor, KGF) regulates lung branching at the distal tips by promoting AEC proliferation (Cardoso et al., 1997; Padela et al., 2008) and FGF10 regulates branching by maintaining cells in a progenitor-like state (Park et al., 1998; Yuan et al., 2018).

To determine the effects of WNT and FGF signaling on organoid growth, we treated AEC-ON 3D cultures with the GSK3 inhibitor, CHIR99021 (CHIR), or a mixture of FGF7 and FGF10 protein ligands (FGF7+10). GSK3 kinase negatively regulates the WNT/β-catenin pathway by maintaining the phosphorylation state of β-catenin (Wu et al., 2010). Inhibition of GSK3 kinase activity results in activation of WNT signaling. **Figure 6A** shows brightfield images of representative wells for vehicle (DMSO), CHIR-, and FGF7+10-treated AEC-ON cells after 2 months of organoid growth. The median sphere sizes for vehicle-treated AEC-ON cells were consistent with median sphere sizes reported under no additive conditions (**Figure 4 – Source Data 1** compared to **Figure 6 – Source Data 1**). Treatment of AEC-ON cultures with either CHIR or FGF7+10 resulted in a statistically significant difference in sphere size. Under CHIR, median sphere size increased from 83 μm to 155 μm (p-value < 2.2 × 10^−16^). Under FGF7+10 treatment, median sphere size increased from 83 μm to 174 μm (p-value < 2.2 × 10^−16^) (**Figure 6B**). Sphere formation efficiencies did not change upon either CHIR or FGF7+10 treatment (**Figure 6C**). AEC-FT and AEC-TN cells were also treated with CHIR or FGF7+10 in 3D culture (**Figure 6 – Supplement 1A, D**). After 2 months of sphere growth, no change in sphere size was detected for either line (**Figure 6 – Supplement 1B, E**). Treatment of AEC-FT cells with FGF7+10 resulted in an increased mean percentage sphere efficiency from 0.3 ± 0.1% to 0.6 ± 0.2% (p-value = 0.03) (**Figure 6 – Supplement 1C**). We noted an increase in number of larger sized “outlier” spheres under FGF7+10 treatment in both AEC-FT- and AEC-TN cultures compared to either vehicle or CHIR conditions (data points lying outside of the upper whisker in **Figure 6 – Supplement 1B, E**). **Figure 6 – Source Data 1** summarizes the growth metrics for the treatment study. In sum, we found that changes in sphere size did not correlate with changes in sphere formation efficiency and that AEC-ON organoids were WNT and FGF responsive, reminiscent of human AEPs.

**Figure 6.**
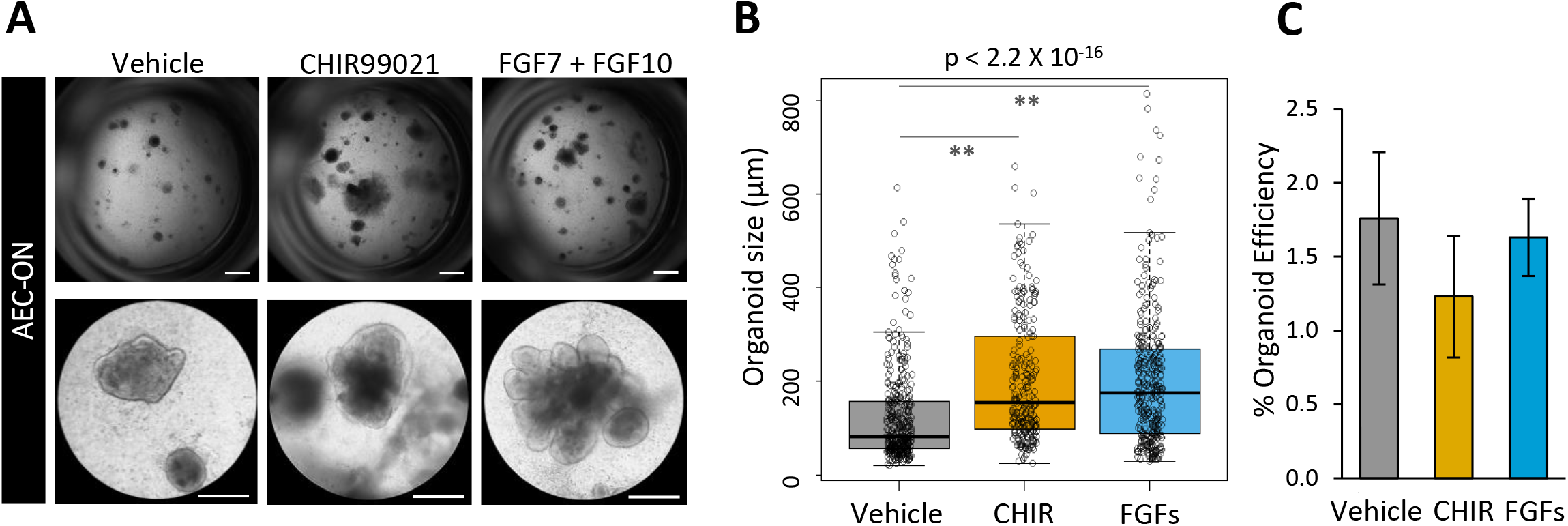
WNT- and FGF treatment of AEC-ON cells in 3D culture results in larger sized organoids. Three-dimensional co-cultures of AEC-ON cells were treated with either vehicle (DMSO), 1 μM Wnt agonist CHIR99021, or a mix of 10 ng/mL FGF7 and 10 ng/mL FGF10 (FGFs) for 2 months. **A)** Whole-well and 10X magnification images of a representative well, per treatment condition. Top scale bar, 1000 μm, bottom scale bar 360 μm. **B)** Organoid size and **C)** organoid formation efficiency were measured for organoids of diameter > 20 μm after 2 months. Plotted values are centered on mean ± standard deviation; n = 4 independent experiments in technical duplicates. *p<0.05, **p<0.005 by nonparametric Wilcoxon test.

## DISCUSSION

The lung is highly susceptible to environmental damage, but is exquisitely engineered to deal with these insults. Upon lung injury, specific cells are mobilized to aid in repair. Depending on the region injured, improper repair and regeneration of the epithelium results in a variety of acute and chronic pulmonary diseases. Having a full spectrum of cell models available to investigate the complete pathophysiology of these diseases would greatly benefit the field. Here, we established a practical method for immortalizing alveolar epithelial cells from human adult lung tissue by first expanding purified AT2 cells in ROCK inhibitor medium followed by SV40 LgT lentiviral transduction. We show that the method works robustly, using cells from three different donor lungs. Gene expression analyses of the cell lines show they are transcriptomically distinct from primary alveolar epithelial cells, lung fibroblasts, and LUAD cancer lines. Our novel AEC-LgT lines proliferate well in 2D culture and despite the absence of mature lung markers, retain the ability to form lung organoids expressing markers of alveolar epithelial cells in 3D co-culture.

Our original goal was to derive immortalized cell lines from purified human AECs. In the process of accomplishing this, we incidentally found that the cells exhibited gene signatures suggestive of either an “intermediate” alveolar epithelial cell state or a distal lung progenitor cell state. In the lung, the alveolar epithelium is maintained by proliferation of AT2 cells and transdifferentiation of a subpopulation of AT2 daughter cells into AT1 cells (Evans et al., 1975; Barkauskas et al., 2013; Nabhan et al., 2018). Liebler *et al*. (2015) found remarkable cellular heterogeneity within the adult lung under homeostatic conditions, identifying populations of ‘intermediate’ or ‘transitory’ AECs expressing combinations of AT2 and AT1 cell markers. AT2 cells in transition (‘late AT2’ cells and ‘early AT1’ cells) lacked SFTPC, but expressed both NKX2-1 and mature AT1 cell markers, AQP5 and HOPX. Interestingly, AQP5 and HOPX did not always overlap. While early *in vitro* transdifferentiation studies did not suggest functions for these intermediate cells, recent studies using scRNA-seq reveal potentially new roles for intermediate or transitory lung cells in injury resolution, which when disrupted may facilitate disease progression, particularly in IPF (Adams et al., 2020; Strunz et al., 2020). AT2 cells have long been considered the cell of origin for many distal lung diseases, but increasing reports from mouse studies are finding that AT1 cells exhibit more cellular plasticity than originally thought; a limited number of AT1 cells is capable of proliferating and transdifferentiating into AT2 cells following pneumonectomy *in vivo* and can generate alveolar-like organoids *in vitro* (Wang et al., 2018; Yang et al., 2016; Jain et al., 2015; Kanagaki et al., 2020). Thus, AT1 cells may not only play a role in alveolar regeneration, but due to their plasticity may also be a source of lung diseases such as cancer or fibrosis. Our characterization studies suggest that cells comprising the AEC-LgT organoids generally represent an intermediate AEC state, exhibiting AT1-like cell expression patterns (AQP5^+^/GPRC5A^+^;SFTPC^−^/HTII280^−^) and possessing AT2-derived ‘alveolosphere’ structural complexity recently reported by Katsura *et al.* (2020).

Distal lung progenitors in the mouse adult lung have been identified as rare subpopulations within the greater ‘bulk’ AT2 population and are not all equally fated. AXIN2^+^/TM4SF1^+^ resident AEPs, comprising ~20% of bulk AT2 cells, give rise to lineage-labeled AT2 and AT1 cells at sites of lung injury after H1N1 influenza infection (Zacharias et al., 2018). AT2 ‘ancillary’ progenitors express AXIN2 when activated upon lung injury, induced by WNT signaling from surrounding stromal cells (Nabhan et al., 2018). While intermediate alveolar cells and alveolar progenitors exhibit different gene expression patterns, the presence of these cell types indicates that the alveolar epithelium has considerable cellular plasticity, which primes the lung to respond to injury expeditiously despite its slow turnover. We discovered our AEC-LgT organoids, particularly AEC-ON cells, exhibit expression patterns of both alveolar ‘intermediate’ and progenitor cells, expressing both NKX2-1 and AQP5, AXIN2, and TM4SF1. Furthermore, AEC-ON organoids were WNT- and FGF-responsive, reminiscent of AEPs, suggesting that under certain growth conditions these cells may be induced to differentiate into mature AT2 or AT1 cell lineages. We are currently investigating this possibility.

In our characterization of the three AEC-LgT cell lines, we found that although these lines were more similar to each other than to primary human AECs or lung fibroblasts, they showed differences in 2D growth kinetics and in the frequency and complexity of organoids formed. These differences could arise from many sources, including genetic differences between subjects, epigenetic differences related to numerous factors including age, gender, environmental exposures, manner of death, and/or ventilation time, and technical differences between experiments related to AEC preparation or cellular response to culture conditions. These aspects will be important to study as additional cell lines are made from a wider range of individuals.

Each of the AEC-LgT cell lines we derived will likely have specific applications of interest for further study. Because the cell lines are easy to expand in 2D culture, they can also be genetically manipulated using genome engineering strategies to develop a series of isogenic cell lines with altered genes of interest, allowing studies of defined differences (including single nucleotide polymorphisms or SNPs) in the same genetic background. In the near future, we intend to apply our methodology to lung tissues from diverse racial/ethnic individuals, filling a longstanding void in research tools to study diseases in underrepresented groups. Our cell lines will likely be easier to use for many laboratories than iPSC-derived AECs, which require extensive stem cell expertise to properly develop and maintain. We envision that human AEC lines will be widely applicable to study the roles of a variety of lung diseases affecting the distal lung, such as cancer, emphysema, and pulmonary viral infections.

## ACKNOWLEDGEMENTS AND FUNDING

## Acknowledgements

The authors wish to thank Monica Flores and Juan Ramon Alvarez for their assistance in isolating and cryopreserving human AT2 cells, members of the Offringa and Marconett laboratories for their helpful critiques, members of the Ryan lab for sharing reagents and imaging resources, Bernadette Masinsin and Jeff Boyd at the USC Broad Stem Cell Core for help with FACS, Yibu Chen and Meng Li of the USC Libraries Bioinformatics Services for assisting with data analyses, and USC’s Center for High-Performance Computing for providing computing resources.

## Funding

This study was supported by NIH grant (1 R01 HL114094) to IAO and ZB, R35 HL135747 to ZB, the Hastings Foundation and generous donations from Conya and Wallace Pembroke and Judy Glick. ET was supported by the California Community Foundation BAPP-15-121814 and a Hastings Center for Pulmonary Research postdoctoral fellowship. ZB is the Ralph Edgington Chair in Medicine. The bioinformatics software and computing resources used in the analysis are funded by the USC Office of Research and the USC Libraries. This work was supported in part by the Norris Comprehensive Cancer Center core grant, award number P30CA014089 from the National Cancer Institute. The content is solely the responsibility of the authors and does not necessarily represent the official views of the National Cancer Institute or the National Institutes of Health. The funders had no role in study design, data collection and analysis, decision to publish, or preparation of the manuscript.

## COMPETING INTERESTS

The authors have no competing interests.

## MATERIALS AND METHODS

### Ethics statement

Remnant human transplant lungs were obtained in compliance with the University of Southern California Institutional Review Board-approved protocols for the use of human source material in research (HS-07-0060). Lungs were processed within 3 days of death. All donors were de-identified. Mouse experiments were performed under the guidance of the University of Southern California Institutional Animal Care and Use Committee (IACUC protocol ID 21116).

### Isolation and culture of primary human alveolar epithelial cells

Lung tissue was collected from de-identified, cancer-free donors: Lung-FT, 25 years old (yr), Caucasian, male; Lung-ON, 66 yr, Caucasian, female; Lung-TN, 62 yr, Caucasian, male (**Table 1**). Human AT2 cells were isolated and purified as previously described (Ballard et al., 2010), modified by using anti-EpCAM beads and purity was assessed by staining of cytospins (**Table 2**). Cells were resuspended in 50:50 growth medium [50% DMEM-F12 (Sigma-Aldrich D64421), 50% DMEM High glucose (Gibco 21063) supplemented with 10% FBS (Omega Scientific FB-11), Pen/Strep, Gentamycin, and Amphotericin B]. The ability to differentiate into AT1-like cells was assessed by *SFTPC* and *AQP5* expression by qPCR and Western blot analyses as described by Marconett *et al.* (2013). Remaining purified AT2 cells were resuspended in 90% growth medium-10% DMSO, frozen in cryovials at 1-2×10^6^ cells/mL, and stored in liquid nitrogen.

**Table 2.** Lung cell purity metrics.

| **Lung Sample** | **Preparation Date**<br>**(days post-tissue approval)** | **SFTPC<sup>+</sup>** | **NKX2-1<sup>+</sup>** | **CD45<sup>+</sup>** | **VIM<sup>+</sup>** | **TUBB4A<sup>+</sup>** |
| --- | --- | --- | --- | --- | --- | --- |
| Lung-FT | 2 | 88% | 96% | 3% | 4% | 3% |
| Lung-ON | 3 | 79% | -- | 15% | -- | -- |
| Lung-TN | 3 | 84% | 86% | 5% | 12% | -- |

Following anti-EpCAM bead purification, cell suspensions were assessed for AT2 cell enrichment by cytopspin preparation, then IF staining with AT2 lung markers (SFTPC, NKX2-1), hematopoietic marker (CD45), fibroblast marker (VIM), and lung airway cell marker (TUBB4A).

### Mycoplasma and rodent pathogens testing

Cells used in the study were negative for mycoplasma and rodent pathogens. Cells were routinely tested using an in-house qPCR-based method adapted from Ishikawa *et al.* (2006). Briefly, cells to be tested were passaged two times in antibiotic- and antimycotic-free media, then collected for genomic DNA (gDNA) extraction using Qiagen DNeasy Blood and Tissue kit (Qiagen 69504) following the manufacturer's instructions with the exception of the last step in which gDNA was eluted in DNase-RNase-free water. Genomic DNA was diluted to a concentration of 10 ng/μL in water, then 50 ng gDNA was used per qPCR reaction using iQ SYBR Green Supermix (Bio-Rad 1708880). Mycoplasma-specific primer sequences are listed on **Table 3**. Each assayed primer set was tested in technical triplicates. Per 50 ng gDNA (5 μL), 0.375 μL 3 μM forward primer, 0.375 μL 3 μM reverse primer, and 6.25 μL SYBR Supermix was combined, mixed gently, then run on an MJ DNA Engine Opticon 2 Research thermocycler. Cycling conditions: Initial 94°C, 3 min, followed by 40 cycles of: (i) 94°C for 15 s; (ii) 65°C for 30 s, (iii) 72°C for 30 s, followed by final extension at 72°C for 10 min. A melting curve (55°C to 95°C) was performed at the end of the PCR to confirm the identity of each product and verify controls. Results were considered negative if Ct values were above 30 cycles. Negative controls typically have Ct between 35-40, positive controls around Ct 15-20. In cases where Ct values were not clear, samples were sent to the Norris Cancer Center Bioreagent and Cell Culture Core for additional testing. For cell line injections into mice, rodent pathogen testing was managed through the USC Department of Animal Resources, which sent cell samples and Matrigel to Charles River Laboratories for testing.

**Table 3.**
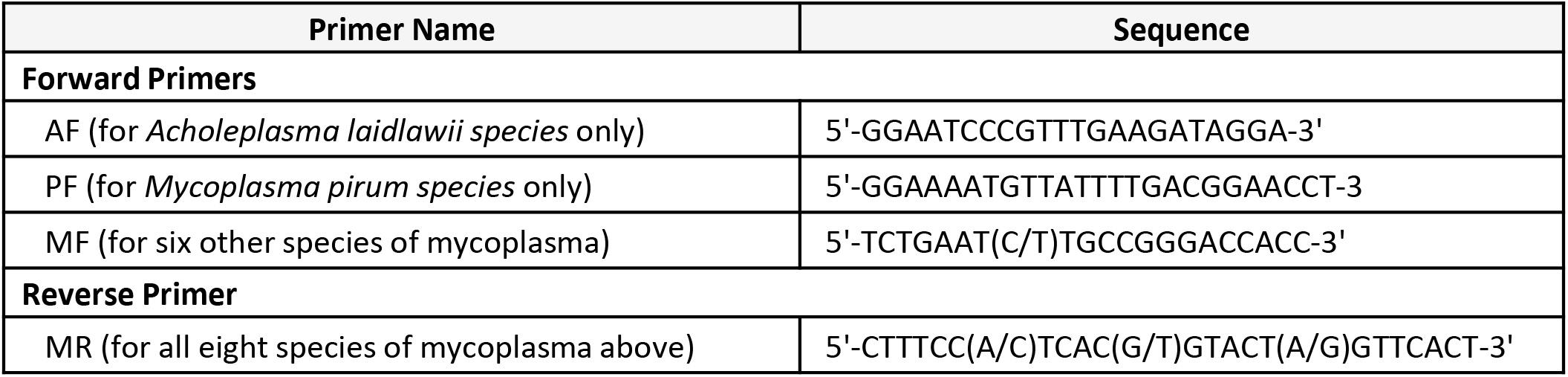
Mycoplasma detection primers.

### Derivation of human alveolar epithelial cell lines

Previously frozen isolated AT2 cells were quick-thawed in a 37°C water bath, spun down to remove freezing medium, and resuspended in Fmed+ROCKinh medium, modified from Liu *et al*. (2012) [3:1 (v/v) DMEM/F12 (Corning 10-090-CV) to DMEM (Gibco 21063-029), 5% FBS, 0.4 μg/mL hydrocortisone (Sigma-Aldrich H0888), 5 μg/mL insulin (Sigma-Aldrich I0516), 8.4 ng/mL cholera toxin (Sigma-Aldrich C8052), 10 ng/mL human recombinant EGF (ThermoFisher PHG0311), Antibiotic-Antimycotic (Gibco 15240-062), and 10 μM Y-27632 (Enzo Life Sciences 270-333)].

Resuspended AT2 cells were plated in 96-well Primaria culture plates (BD Falcon 3872), ~6250 cells per well and allowed to attach for 2 days. On the second day, Fmed+ROCKinh medium was completely replaced with fresh medium. Wells were monitored every day for surviving cells and proliferation. Media were changed every 2-3 days. Once cells reached 90-100% confluence, cells were detached with Accutase (Innovative Cell Technologies AT-104) and re-plated onto 48-well culture plates (“Passage 1”). This procedure was performed again, re-plating cells onto 24-well culture plates (“Passage 2”), then 12-well culture plates (“Passage 3"), then 6-well culture plates (“Passage 4”). Stocks of cells were frozen down starting at passages 3 and 4 in medium containing 10% DMSO and 90% 0.22 μm filtered FBS.

### Construction of lentiviral plasmids and production of viral particles

LeGO iG and LeGO iT plasmids (http://www.lentigo-vectors.de/) were kind gifts from Dr. Kate Lawrenson (Cedars Sinai Medical Center, Los Angeles, CA). CDK4^R24C^ coding sequence (CDS) was subcloned from pBABE-hygro-CDK4^R24C^ plasmid (Addgene 11254) by PCR amplification into LeGO iG vector between *Bam*HI and *Sbf*I sites, upstream of enhanced green fluorescent protein (eGFP). hTERT CDS was subcloned from pBABE-puro-hTERT plasmid (Addgene 1771) into LeGO iT vector between *Bgl*II and *Eco*RI sites, upstream of tdTomato. SV40 Large T antigen CDS was subcloned from pBABE-puro-SV40 LgT plasmid (Addgene 13970) into LeGO iG vector between *Bam*HI and *Eco*RI sites, upstream of eGFP. Plasmids were propagated in Stbl3 chemically competent *E.coli* (ThermoFisher C737303). Plasmid sequences were verified by Sanger sequencing (GENEWIZ Inc).

Third generation lentiviral particles were produced in low passage 293T cells by transfection. Per 10 cm dish of 4-5×10^6^ 293T cells, we used: 15 μL BioT (Bioland Scientific LLC B01-00), 2 μg pCMV-VSVG (Addgene 8454), 2 μg pMDLg/pRRE (Addgene 12251), 2 μg pRSV-Rev (Addgene 12253), and 2 μg lentiviral plasmid carrying transgene (LeGO iG-CDK4^R24C^, LeGO iT-hTERT, LeGO iG-SV40 LgT). Viral supernatant was collected at 48- and 72-hours post-transfection, pooled, spun down at 300 g to remove cell debris, filtered through 0.45 μm PES filters, and concentrated using Lenti-X Concentrator (Takara Bio 631231). Lentiviral pellets were resuspended in DMEM, aliquoted, and stored at −80°C. Transduction efficiency was tested empirically on 293T cells.

### Lentiviral transduction of human AECs

AEC-ROCKinh cells (passage 4) were plated onto 96-well culture plate in Fmed+ROCKinh media. The following day when cells were at 40-50% confluence, different volumes of the lentiviruses, either singly or in combination, were mixed with 8 μg/mL polybrene in Fmed+ROCKinh media and added to each well. Of 300 μL resuspended viral supernatant, 1, 2, and 3 μL of LeGO iG-CDK4^R24C^, LeGO iT-hTERT, and LeGO iG-SV40 LgT viruses were used to transduce cells in a total media volume of 50 μL. The following day, 100 μL of Fmed+ROCKinh media were added to each well to dilute out the viral supernatant. Two days post-transduction when cells were 90-100% confluent, cells were detached using Accutase and re-plated onto 48-well culture plates. At four days post-transduction, expression of CDK4^R24C^, hTERT, and SV40 LgT was checked by fluorescence microscopy (Nikon Eclipse Ti-U inverted fluorescence microscope). Cells transduced with LeGO iT-hTERT only, LeGO iT-hTERT + LeGO iG-CDK4^R24C^, or LeGO iT-hTERT + LeGO iG-SV40 LgT were sorted by fluorescence-activated cell sorting (FACS) on eGFP, tdTomato, or dual fluorescence at the USC Flow Cytometry Core Facility (FACS Aria II, BD Biosciences). Negative fluorescence was set by AEC-ROCKinh cells. All cell lines were maintained in Fmed+ROCKinh media.

### Proliferation assay

One thousand cells were plated on 24-well culture plates in quadruplicate and monitored for seven days. Twenty-four hours post seeding (day 1), cells were detached with Trypsin-EDTA (0.05% Trypsin, 0.02% EDTA) and resuspended in growth media. Cells were counted manually using a hemocytometer. Cell counts were reported as average total cell number ± standard deviation from at least three biological replicates. Population doubling time (PDT) was calculated based on the linear part of the growth curve using the equation [(t_2_-t_1_)/3.32] X (log n_2_ – log n_1_), where n_2_ was the number of cells on day 6 and n_1_ was the number of cells on day 4. High density proliferation assays were performed as described above with an initial cell seed count of 5000 cells per well.

### Anchorage-independent growth assay

Per well of a 6-well culture plate, 1.5 mL of 0.6% (w/v) Difco Noble Agar (BD Biosciences 214220) in Fmed+ROCKinh media was added to form the bottom layer of the soft agar assay. For the top layer, in 1.5 mL, five thousand cells were mixed with 0.3% final concentration Noble Agar in Fmed+ROCKinh media. The top layer was allowed to solidify at room temperature (RT) before 1 mL Fmed+ROCKinh media was carefully added. For A549 positive control cells (ATCC CCL-185), RPMI 10% FBS was used to set up soft agar layers. Media for all cell lines were changed every 3 days. Colony growth was monitored for 1 month. Colonies were visualized by staining soft agar samples with crystal violet solution (crystal violet dissolved in 10% ethanol) according to Borowicz *et al.* (2014) and counted using ImageJ software. Experiments were replicated biologically at least three times, each with six technical replicates. Data were reported as mean number of colonies ± standard deviation.

### Subcutaneous injection of nude mice

Tumorigenicity of the three AEC-LgT cell lines was assessed by subcutaneous injection of 1×10^6^ cells into each flank of 6 week old, male and female homozygous Foxn1^nu^, NU/J mice (Jackson Laboratories, strain 002019). NU/J mice were kept in sterile housing and given irradiated rodent feed *ad libitum*. On the day of injection, cells were detached with Accutase, collected in Fmed+ROCKinh growth media, and spun down. Supernatant was removed, cells were resuspended in 1 mL growth media, and manually counted using a hemocytometer. For each flank, a master mix of cells resuspended in sterile phosphate-buffered saline (PBS) and 50% final concentration of Matrigel Membrane Matrix solution (Corning 354234) was prepared such that 1×10^6^ cells were delivered in 150 μL of solution. Mixes were kept on ice as injections were performed to prevent premature solidification. Cell mixtures were injected using a tuberculin syringe with attached 27G needle (BD 305620). Mice were anesthetized by inhaled isoflurane according to IACUC-approved procedures. AEC-hTERT cells were used as negative controls; A549 cells were used as positive controls. We used 4 mice for A549 cells and 8 mice for each AEC line, with equal numbers of males and females. Injections were performed as a single-blind study, where sample identities were unknown to the experimenter performing injections. Mice were monitored every 3 days, weighed every week, and nodule length (L) and width (W) were measured with a digital caliper (iKKEGOL model 714838802360, Shenzen, China) weekly starting 3 weeks post-injection. Nodule volume (V) was calculated as V = L X (W/2)^2^ (Luo et al., 2018). Excised nodules were fixed in 4% paraformaldehyde (PFA) overnight at 4°C, processed for paraffin embedding and sectioning by the Norris Cancer Center Pathology Core at USC. An expert lung pathologist blind to sample identities was consulted to evaluate H&E nodule sections.

### Three-dimensional (3D) co-culture

Actively dividing AEC-LgT cells between passages 6 and 19 were used to set up 3D co-culture with neonatal mouse lung fibroblasts, MLg (ATCC CCL-206). MLgs were cultured in EMEM 10% FBS and maintained at sub-confluence. Noticeably lower organoid formation efficiency was observed when MLgs that had been grown beyond 70% confluence were used in the 3D co-culture. Five thousand AEC-LgT cells were mixed with 50,000 MLgs in Basic medium [phenol-red free DMEM/F12 (Gibco 11039021), 1X ITS (Gibco 41400-045), 10% FBS, 1X Antibiotic-Antimycotic] with 50% Growth Factor Reduced Matrigel (Corning 354230) and plated on Clear Transwell inserts (Corning 3470), 100 μL per insert. Basic medium supplemented with 10 μM SB-431542 (BioVision 1674), a transforming growth factor beta (TGFβ) inhibitor, was added to the outer chamber and replaced every 2 days. Organoid formation was monitored under brightfield microscopy for 1-2 months. Whole-well images were captured using a Leica MZ16 F fluorescence stereomicroscope and Spot Advanced software (v4.5.8) through the USC Hastings Center for Pulmonary Research Core. Brightfield and fluorescence images at 4X magnification were captured using ECHO Revolve R4 fluorescence microscope (San Diego, CA). Organoid size in microns was measured using the “Annotation length” feature on the ECHO Revolve R4 microscope; reported sizes are based on the longest diameter. Organoid number was determined using the “Annotation count” feature on ECHO Revolve R4 microscope. Values were reported as the mean ± standard deviation.

### Treatment of 3D co-cultures

Three-dimensional co-cultures were set up as described above. Treatment with 1 μM WNT agonist CHIR99021 (Sigma-Aldrich SML1046) dissolved in DMSO or 10 ng/mL FGF7 (Peprotech 100-19) and 10 ng/mL FGF10 (Peprotech 100-26) dissolved in sterile PBS began two days following culture set up, where SB Basic medium was replaced with fresh SB media containing either CHIR99021 or FGF7+FGF10, or DMSO (vehicle control). Media were changed every two days. Cultures were maintained for two months before sphere size and number were assessed.

### Histological processing of organoids

Once organoids formed, inserts were removed and fixed with 4% PFA for 30 min. PFA solution was removed by inverting inserts. Inserts were then submerged in 1X PBS for 15 min with two changes and dehydrated in 70% ethanol for 30 min with three changes. Matrigel samples were removed from insert housing by cutting the filter out from the bottom face with a feather razor, then embedded in Histogel (ThermoFisher HG-4000) and equilibrated in 70% ethanol for 1 h at RT. Histogel “buttons” were further dehydrated using standard methods, then embedded in paraffin wax. Paraffin blocks were sliced to 5 μm sections in-house using Microm HM 314 microtome through the USC Hastings Center for Pulmonary Research Core.

### Immunofluorescence staining

#### For 2D cultures

Cells were plated on standard multiwell culture plate to reach confluence the following day. Cells were rinsed with filtered 1X PBS, fixed with ice cold methanol for 10 min, washed three times with PBS, blocked with 5% filtered bovine serum albumin (BSA) in PBS, then probed overnight at 4°C with respective primary antibodies in 5% BSA-PBS solution. Horse serum (RMBIO DES-BBT) diluted to 30% in PBS was used as blocking solution for pro-SPC antibody. The following day, cells were washed with 1X TBST (20 mM Tris, 150 mM NaCl, 0.01% Tween 20, pH 7.5), probed in PBS with biotinylated secondary antibodies for 1 h, washed, then probed with Streptavidin-Alexa Fluor 647 conjugate (ThermoFisher S21374). For double staining, cells were blocked again, then probed with either donkey anti-rabbit Alexa Fluor 488 secondary antibody (ThermoFisher A21206) or donkey anti-mouse Alexa Fluor 488 secondary antibody (ThermoFisher A21202). Mounting solution with 4′,6-diamidino-2-phenylindole (DAPI) was used as nuclear counterstain (Vector Laboratories H-1200). Stained cells were viewed using Nikon Eclipse Ti-U inverted fluorescence microscope and imaging software, NIS-Elements Br (v4.00.12, build 802, 64-bit).

#### For 3D cultures

Organoid paraffin sections were baked in a 60°C oven for 12 hours. Excess paraffin was wiped off with Kimwipes. Slides were then submerged in xylene, two changes, 5 min each, rehydrated through a series of ethanol baths, each with two changes (100% ethanol, 95%, 85%, 75%, 50%), and rinsed with distilled water. Samples were boiled for 6 min on high power, then for 5 min at 10% power in Tris-based antigen unmasking solution (Vector Laboratories H-3301) in a standard microwave oven, cooled to RT, then permeabilized with 2% Triton X-100 in PBS for 15 min, washed with PBS, then blocked with 5% BSA-PBS or 30% horse serum in PBS (for pro-SPC antibody) for 1 h RT. Citrate-based unmasking solution was used for pro-SPC probed samples (Vector Laboratories H-3300). Primary antibodies were diluted in 5% BSA-PBS or 30% horse serum (pro-SPC antibody) and probed overnight at 4°C. Subsequently, all washes were with 1X TBST. For single and double stainings, biotinylated secondary antibodies, fluorochrome-conjugated secondaries, Streptavidin-Alexa Fluor 647, Streptavidin-Alexa Fluor 488 (ThermoFisher S11223), and Streptavidin-FITC antibodies (ThermoFisher SA-10002) in PBS were incubated 1 h at RT for each step. Sections were mounted with Prolong Gold antifade reagent with DAPI (Invitrogen P36931), sealed, cured overnight at RT, then stored at 4°C. Slides were visualized the following day using the ECHO Revolve R4 fluorescence microscope.

#### Antibodies

Primary antibodies were: HOPX (SCBT sc-30216), pro-SPC (Seven Hills Bioreagent WRAB-9337), SV40 LgT (SCBT sc-147), AQP5 (Abcam ab92320), ECAD (BD Transduction Laboratories 610181), NKX2-1 (Leica TTF-1-L-CE), Ki67 (Abcam ab16667), SOX2 (SCBT sc-365823), SOX9 (SCBT sc-20095), GPRC5A (Abbexa ab005719), HTII280 (Terrace Biotech TB-27AHT2-280), KRT5 (Abcam ab52635), mouse control IgG (Vector Laboratories I-2000), rabbit control IgG (Vector Laboratories I-1000), mouse control IgM (Sigma-Aldrich M5909). Secondary antibodies were: biotinylated horse anti-mouse IgG (Vector Laboratories BA-2000), biotinylated goat anti-rabbit IgG (Vector Laboratories BA-1000), biotinylated goat anti-mouse IgM (Vector Laboratories BA-2020). See **Table 4. Key Resources Table** for additional reagent details.

**Table 4.**
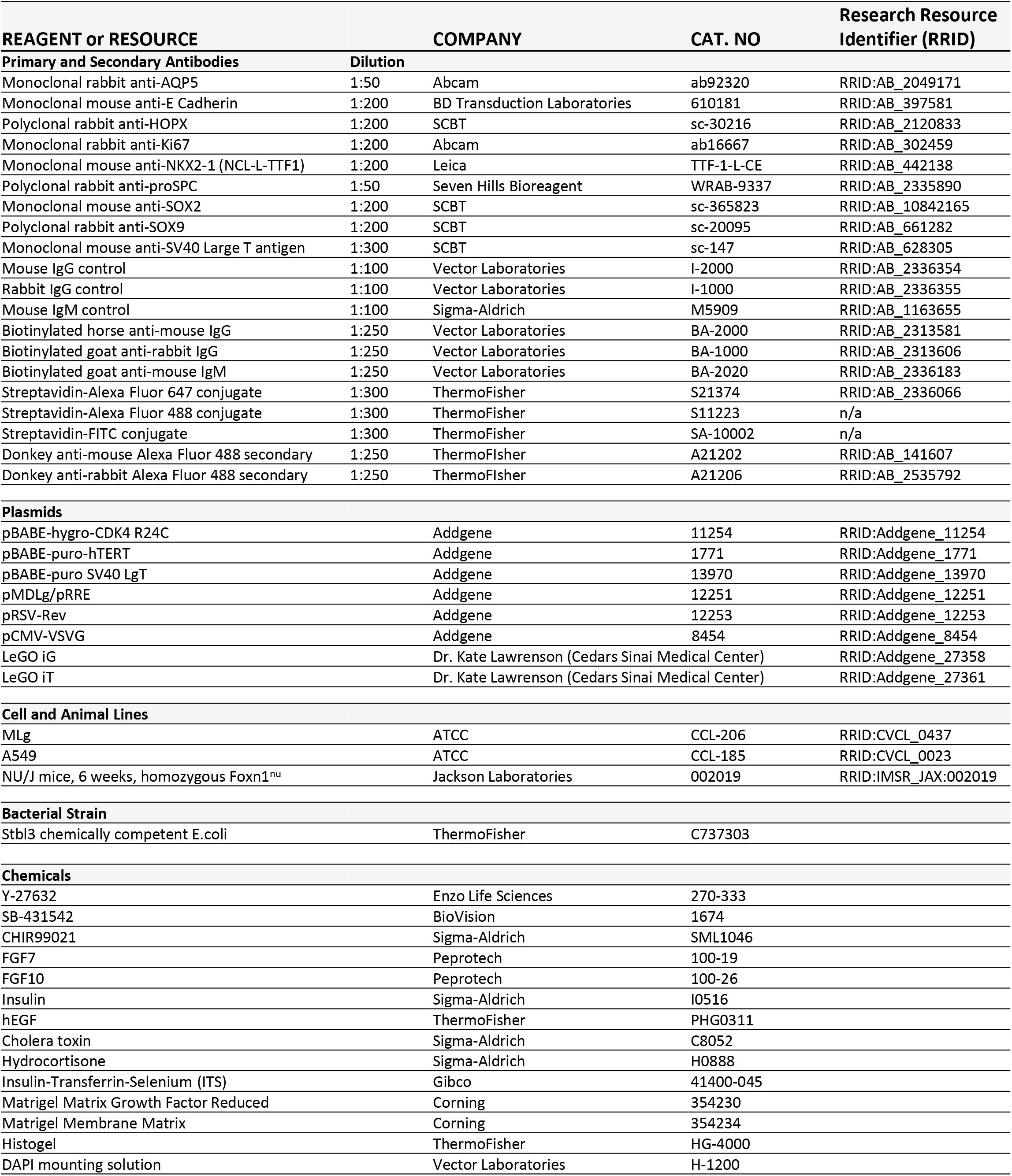
Key Resources Table.

### Statistical Analyses

The Wilcoxon nonparametric rank sum test on independent samples was performed for anchorage-independent growth assays, 3D organoid size, and percent organoid forming efficiency comparisons. Wilcoxon tests were performed using the R platform (v3.6.0 “Planting of a Tree”) with the function *wilcox.test()* on the appropriate datasets. The student’s t-test was performed for proliferation assays using the R software function *t.test()* (v3.6.0).

### Bulk RNA-sequencing analyses

RNA-seq analyses comparing derived AEC lines with normal lung cells and LUAD cancer cell lines were performed using publicly available data from ENCODE (https://www.encodeproject.org/) and DBTSS (https://dbtss.hgc.jp/) databases (**Figure 2 – Source Data 1**). For the AEC lines (AEC-FT-ROCKinh, AEC-CDK4^R24C^, AEC-hTERT, AEC-CDK4^R24C^+hTERT, and AEC-FT) and the adult human lung fibroblast cell line, HLF-133, total RNA was isolated from subconfluent, exponentially dividing cells using Illustra TriplePrep kit (GE Healthcare 28-9425-44) following manufacturer’s instructions. For the primary AECs, purified alveolar epithelial cells from three de-identified human donor lungs were used. Cells from each lung were distinguished from each other by the sample name extension “-m,” “-f,” or “-a.” From each lung, purified primary AT2 cells and AT2 cells transdifferentiated on filters for 6 days into AT1-like cells (Danto et al., 1995) were isolated and total RNA extracted. Total RNA from purified AT2 cells and AT1-like cells from one of the donor lungs (labeled “AEC-m” in **Figure 2 – Source Data 1**) had been previously characterized (Yang et al., 2018; GEO record GSE84273). Total RNA from the remaining two donor lungs (“AEC-f” and “AEC-a” in **Figure 2 – Source Data 1**) was also previously isolated in our laboratory and sequenced, however without subsequent publication. For the AEC lines, HLF-133 cell line, and primary AECs, 2 μg of RNA were submitted to the USC Molecular Genomics Core facility for sequencing. RNA quality was assessed on the Bioanalyzer (Agilent) then rRNA-depleted using Ribo-Zero rRNA Removal kit for human samples (Illumina MRZH11124) before proceeding with library preparation (TruSeq mRNA Stranded Library preparation kit, Illumina 20020594). These samples were sequenced paired-end 75 bp (PE75) at a depth of ~ 20-30 million reads per sample, on HiSeq2000/2500 (Illumina). For AEC-ON and AEC-TN cell lines, total RNA was isolated using the Illustra TriplePrep kit as detailed above and 1 μg of RNA was submitted to the UCLA Technology Center for Genomics and Bioinformatics for sequencing. Samples were rRNA depleted and libraries were prepared at the UCLA facility, PE75, sequenced at a depth of ~ 30 million reads per sample on NextSeq500 Mid Output (Illumina). Raw fastq files were retrieved and processed as follows:

Raw fastq files generated from our samples and taken from ENCODE and DBTSS databases were uploaded to Partek Flow through the USC Norris Medical Library Bioinformatics Core and the USC High-Performance Computing nodes. Files were quality controlled using Partek’s QC tool and trimmed at both ends using Partek default parameters, then aligned using STAR RNA-sequence aligner (v 2.6.1d) to the human genome assembly, hg38 GENCODE Genes, release 29. Raw read counts were generated by quantification to the transcriptome using Partek E/M algorithm under default parameters, using hg38 GENCODE, release 29. Raw counts were then analyzed and processed in R (v3.6.0) using the DESeq2 package (Love et al., 2014).

#### Gene clusters from unsupervised clustering

GO terms were analyzed using PANTHER14.1 (2018_04 release) PANTHER Overrepresentation Test (Released 20200728) with the default reference gene list of all *Homo sapiens* genes in the GO Ontology database (released 2019-07-03) (www.geneontology.org). Statistically significant GO terms were calculated using Fisher’s Exact Test with FDR corrected p-value cutoff of < 0.05.

#### GO terms of differentially expressed genes

Associated GO terms for each gene were taken from Ensembl gene database of annotated genes (release 98, Human genome assembly GRCh38.p13, www.ensembl.org).

### Single cell-sequencing analyses

AEC-ON cells grown in 2D and as organoids in 3D co-culture were harvested in parallel by sequential Dispase protease treatment (StemCell Technologies 07923) and gentle centrifugation. Dissociation of organoids into a single-cell suspension was assessed by brightfield microscopy using a hemocytometer. Surrounding single cells in the 3D culture that did not form organoids were excluded from collection by first incubating samples with Dispase briefly for ~15 min in a 37°C water bath, followed by gentle centrifugation. The subsequent iterations of Dispase treatment were for 30 min in a 37°C water bath, followed by centrifugation. Cells were immediately FACS sorted for GFP^+^ epithelial cells using MLg fibroblasts as a negative control for GFP fluorescence gating. Retrieved cells for each sample were then washed in 0.04% BSA-PBS solution. Barcoding and library preparation for scRNA-seq was performed following the manufacturer’s protocol for the 10x Genomics Chromium Single Cell 3’ GEM, Library & Gel Bead Kit v3 (PN-1000092). cDNA libraries were assessed for quality and quantification according to the Chromium Single Cell 3’ Reagent kit v3 user guide using the High Sensitivity D5000 ScreenTape (Agilent 5067-5592) and 2200 TapeStation Controller software (Agilent, Santa Clara, CA). Sequencing was performed using Illumina HiSeq 3000/4000 kit at a coverage of 50,000 raw reads per cell (Paired-end; Read 1: 28 cycles, i7 Index: 8 cycles, i5 Index: 0 cycles, Read 2: 91 cycles). Raw data were processed using the CellRanger function *cellranger count* (10x Genomics, v3.1.0, default settings) to align to the human reference genome (hg19, 10x Genomics, v1.2.0) and identify 4320 2D cells and 7144 3D cells (11,464 total cells), which were then aggregated using the CellRanger pipeline (10x Genomics, v3.1.0, *cellranger aggr* function on default settings). Pre-processed outputs were then analyzed in R using the Seurat package for additional quality control assessment and downstream analyses (Butler et al., 2018; Stuart et al., 2019). Cells with less than 500 transcripts profiled and more than 18% of their transcriptome of mitochondrial origin were removed, leaving a total of 10,965 cells (4082 2D and 6883 3D cells) used for clustering and visualization. Read counts were normalized using the SCTransform method (Hafemeister and Satija, 2019). Dimensionality reduction and clustering analyses were performed as outlined in the Seurat vignette (https://satijalab.org/seurat/v3.2/pbmc3k_tutorial.html) with modifications: a Shared Nearest Neighbor (SNN) graph was constructed using the *FindNeighbors()* function on 20 dimensions of reduction and clusters were determined using the *FindClusters()* function with a reduction of 0.3.

## Notes

### Competing Interest Statement

The authors have declared no competing interest.

