## Supplementary Figures and Tables for "Development of novel *in vitro* human alveolar epithelial cell models to study distal lung biology and disease"

Evelyn Tran[1,2,3], Tuo Shi[1,2,3], Xiuwen Li[2,4], Adnan Y. Chowdhury[5], Du Jiang[5], Yixin Liu[6], Hongjun Wang[6], Chunli Yan[1,2], William D. Wallace[7], Rong Lu[5], Amy L. Ryan[5,6], Crystal N. Marconett[1,2,3], Beiyun Zhou[2,6], Zea Borok[2,3,6], Ite A. Offringa[1,2,3,†]

[1]Department of Surgery, Keck School of Medicine, USC, Los Angeles, CA, USA; [2]USC Norris Comprehensive Cancer Center, Keck School of Medicine, USC, Los Angeles, CA, USA; [3]Department of Biochemistry and Molecular Medicine, Keck School of Medicine, USC, Los Angeles, CA, USA; [4]Department of Translational Genomics, Keck School of Medicine, USC, Los Angeles, CA, USA; [5]Department of Stem Cell Biology and Regenerative Medicine, Eli and Edythe Broad CIRM Center for Regenerative Medicine and Stem Cell Research, Keck School of Medicine, USC, Los Angeles, CA, USA; [6]Hastings Center for Pulmonary Research and Division of Pulmonary, Critical Care and Sleep Medicine, Department of Medicine, Keck School of Medicine, USC, Los Angeles, CA, USA; [7]Department of Pathology, Keck School of Medicine, USC, Los Angeles, CA, USA

### Fig 1 – Supplement 1. Lentiviral constructs used to derive AEC lines

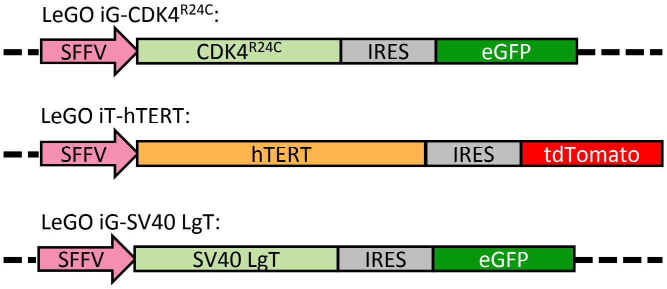


**Figure 1 - Supplement 1. Lentiviral constructs used to derive human alveolar epithelial cell lines.** Diagrams of immortalizing lentiviral constructs carrying, respectively, p16-insensitive $CDK4^{R24C}$ mutant subcloned into the LeGO iG vector with eGFP fluorescence, hTERT subcloned into the LeGO iT vector with tdTomato fluorescence, and SV40 LgT subcloned into the LeGO iG vector with eGFP fluorescence.

### Fig 1 – Supplement 2. Optimization of human AEC line transduction conditions

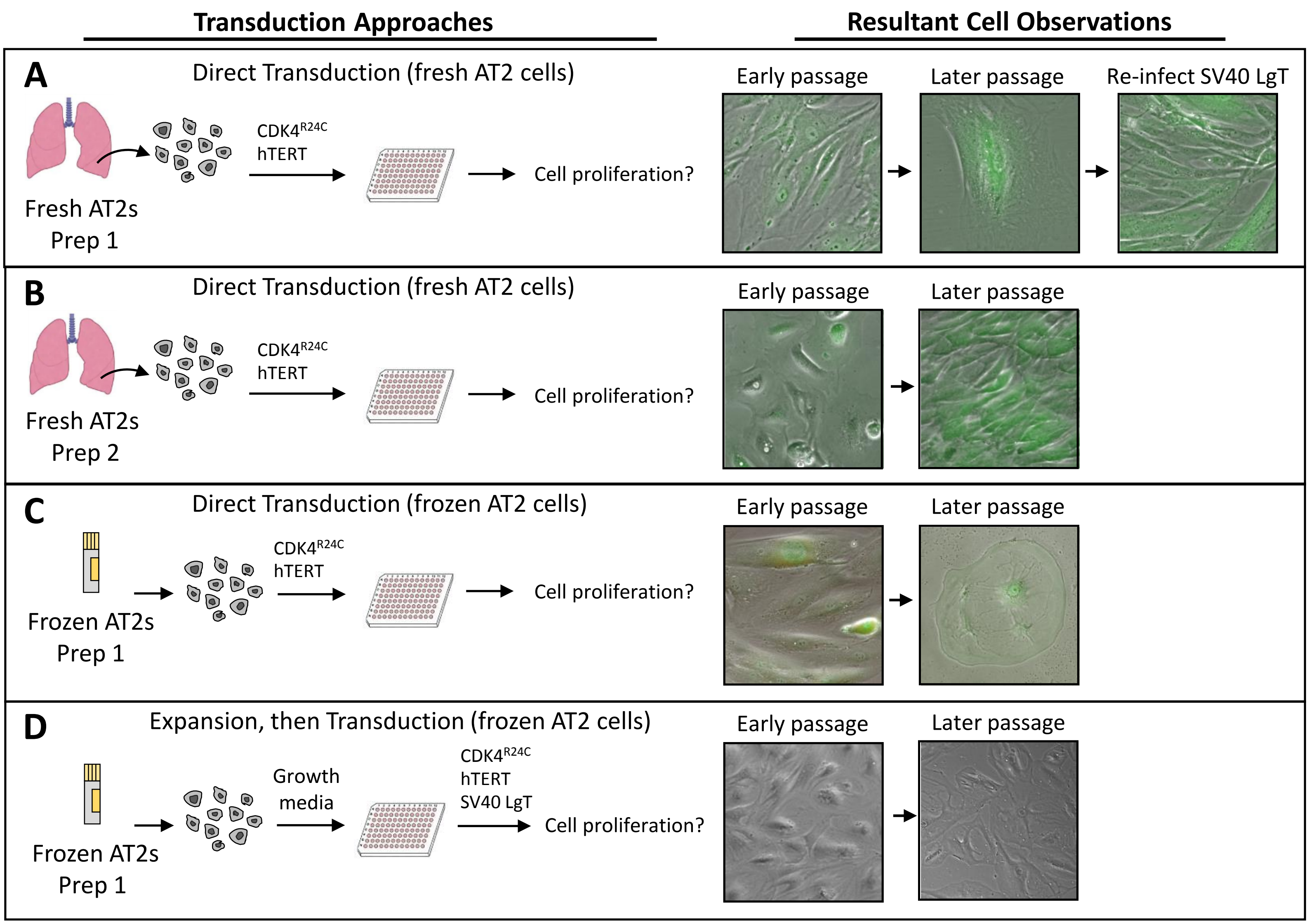


**Figure 1 – Supplement 2. Optimization of immortalized human alveolar epithelial cell culture conditions.** Different strategies for cell immortalization were used on two different preparations (prep) of purified human AT2 cells from two different lungs. **Left images:** Diagram of transduction approaches used to culture primary AT2 cells. "Direct transduction:" Cells were mixed in suspension with different combinations of lentiviruses, then plated into 96-well culture dishes. "Expansion, then transduction:" Cells were plated first in media promoting cell proliferation, then expanded, counted, and plated into 96-well dishes to be transduced with immortalizing lentiviruses. **Right images:** Merged fluorescence or brightfield images of cell morphologies resulting from the different immortalization strategies. "Early passage" refers cells that were passaged fewer than 3 times. "Later passage" refers to cells beyond 3 passages where continued cell division was no longer observable. **A and B)** Direct transduction of freshly isolated AT2 cells. **C)** Direct transduction of cryopreserved AT2 cells from Prep 1. **D)** Cryopreserved AT2 cells from Prep 1 were plated in media and split several times before transduction with lentiviruses. For this study, condition (D) was pursued to derive all AEC lines. Images were taken at 10X magnification.

### Fig 1 – Source Data 1. Media tested in small-scale screen

| | Molecule of Interest | Mechanism of Action | Reported Function | Media conditions | References |
|---|---|---|---|---|---|
| 1 | FBS<br>Fetal bovine serum<br>(No targeted molecule) | N/A | FBS is a common undefined nutrient additive in mammalian cell culture systems since the 1990s.<br>Maintains isolated alveolar epithelial cells in culture | 50/50 DMEM/F12<br>20% heat-inactivated FBS<br>1% Pen/Strep<br>1% Amphotericin B (antimycotic) | "Animal Sera." *ATCC Animal Cell Culture Guide*. <www.attc.org/en/Guides/aspx><br>Fang et al. *Plos One*. 2017; 12(6): e0178960.<br>Borok et al. *Am J Physio* l. 1996 Apr; 270(4 Pt 1):L559-65 |
| 2 | BIO<br>6-bromoindirubin-3'-oxime | ATP-competitive inhibitor of the Ser/Thr kinase, GSKβ; binds to the ATP binding pocket preventing GSKβ activation, β-catenin is stabilized, thereby promoting transcription | Activates Wnt signaling increasing cell proliferation and differentiation<br>Regulates epithelial stem cell homeostasis<br>Maintains undifferentiated phenotype of human and mouse embryonic stem cells by sustaining expression of pluripotent transcription factors | 50/50 DMEM/F12<br>0.1% heat-inactivated FBS<br>1 μM, 2.5 μM , or 5.0 μM BIO<br>1% Pen/Strep<br>1% Amphotericin B (antimycotic) | Fevr et al. *Mol Cell Biol*. 2007 Nov; 27(21):7551-9.<br>Yang et al. *Stem Cells.* 2015 May; 33(5):1670-81.<br>Liu et al. *Am J Respir Cell Mol Biol*. 2015 Jul; 53(1):113-24.<br>Sato et al. *Nat Med.* 2004 Jan; 10(1):55-63. |
| 3 | KGF (FGF7)<br>Keratinocyte Growth Factor | Ligand specifically binds to FGFR2b, causes receptor dimerization, then transphosphorylation of intracellular kinase domains and activation of downstream pathways, RAS-RAF-MAPK, PI3K-AKT | Mitogenic effects on AT2 cells, inhibits apoptosis<br>Promotes migration and wound repair<br>Promotes rat AT2 cell proliferation and maintains AT2 cell phenotype | 50/50 DMEM/F12<br>2% heat-inactivated FBS<br>1% Insulin-Transferrin-Selenium solution<br>100 ng/mL hEGF<br>125 pg/mL, 250 pg/mL, or 500 pg/mL KGF<br>1% Pen/Strep<br>1% Amphotericin B (antimycotic) | Shyamsunder et al. *Am J Respir Crit Care Med*. 2014 Jun; 189(12):1520-9.<br>Qiao et al. *Am J Respir Cell Mol Biol.* 2008 Feb; 38(2):239-46.<br>Borok et al. *Am J Physiol*. 1998 Jan; 274(1):L149-58.<br>Yano et al. *Am J Physiol Lung Cell Mol Physiol*. 2000 Dec; 279(6):L1146-58.<br>Mason et al. *Am J Physiol Lung Cell Mol Physio* l. 2002; 282(2):L249-L258. |
| 4 | Keratinocyte Growth Medium<br>(Lonza, #CC-4131) | Designed to support both clonal growth and high density keratinocyte proliferation while maintaining cell morphology. | Used to maintain primary human epidermal cells in culture in the absence of serum and feeder cells. | DMEM base media + KGM SingleQuots:<br>Bovine Pituitary Extract<br>hEGF<br>Insulin<br>Hydrocortisone<br>1% Pen/Strep | Lamb and Ambler. *PLoS One*. 2013; 8(1): e52494.<br>Costea et al. *J Invest Dermatol*. 2003 Dec; 121(6):1479-86. |
| 5 | Y-27632<br>ROCK inhibitor | Small molecule inhibitor of ROCK1 and ROCK2 Rho-kinases, competes with ATP for binding of ATP binding pocket on ROCK proteins, preventing relief of kinase domain autoinhibition | Promotes proliferation of primary epithelial cells<br>Suppresses dissociation-induced apoptosis in stem cells<br>Enhances recovery and growth of stem cells | (3:1, vol/vol) DMEM/F12:DMEM<br>5% FBS<br>0.4 μg/mL Hydrocrotisone<br>5 μg/mL Insulin<br>8.4 ng/mL cholera toxin<br>10 ng/mL hEGF<br>5 μM or 10 μM Y-27632<br>1% antibiotic-antimycotic | Liu et al. *Am J Pathol*. 2012 Feb; 180(2): 599–607.<br>Claasen et al. *Mol Reprod Dev*. 2009 Aug; 76(8): 722–732.<br>Watanabe et al. *Nat Biotechnol*. 2007 Jun; 25(6):681-6.<br>Zhang et al. *PLoS One*. 2011; 6(3): e18271. |

### Fig 1 – Supplemental 3. High density proliferation assays

A

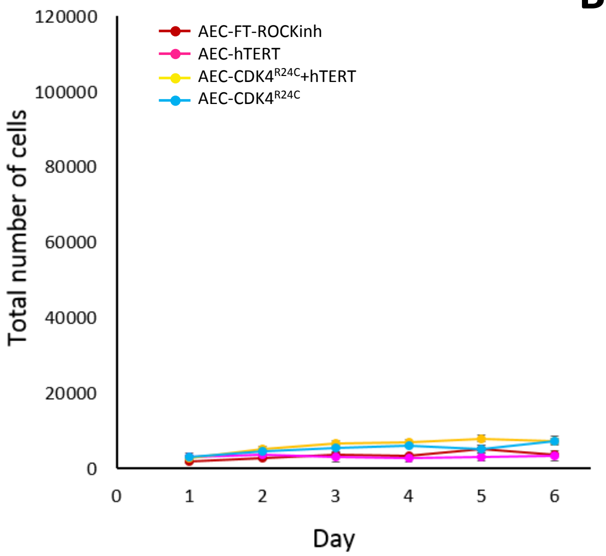


B

| Cell Line | **PDT (mean ± stdev)** (1000/well) | **PDT (mean ± stdev)** (5000/well) |
|---|---|---|
| A549 | 1.3 ± 0.4 days | -- |
| AEC-FT-ROCKinh | no exponential growth | no exponential growth |
| AEC-CDK4$^{R24C}$ | no exponential growth | no exponential growth |
| AEC-hTERT | no exponential growth | no exponential growth |
| AEC-CDK4$^{R24C}$+hTERT | no exponential growth | no exponential growth |
| AEC-FT | 1.1 ± 0.2 days | -- |
| AEC-ON | no exponential growth | 1.1 ± 0.05 days |
| AEC-TN | no exponential growth | 2.1 ± 0.4 days |

**Figure 1 – Supplement 3. High density proliferation assays on slow growing AEC lines. A)** Five thousand cells were seeded per well of a 24-well culture plate and counted every day for 6 days. Total cell numbers were plotted as the mean ± standard deviation, n = 3 independent experiments with technical quadruplets. **B)** Summary table of PDTs for all AEC lines under standard proliferation assay conditions (1000 cells/well) and high density conditions (5000 cells/well). PDTs were only calculated for cell lines that exhibited exponential growth. Also see Figure 1D and E.

### Fig 2 – Source Data 1. List of data samples and sources

| SAMPLE NAME | SAMPLE TYPE | SOURCE | ID* | ACCESSION NUMBER** | SOURCE WEBSITE |
|---|---|---|---|---|---|
| AEC-FT_rep1 | AEC line | IAO | | | |
| AEC-FT_rep2 | AEC line | IAO | | | |
| AEC-hTERT | AEC line | IAO | | | |
| AEC-CDK4$^{R24C}$ | AEC line | IAO | | | |
| AEC-CDK4$^{R24C}$+hTERT | AEC line | IAO | | | |
| AEC-FT-ROCKinh | AEC line | IAO | | | |
| AEC-ON | AEC line | IAO | | | |
| AEC-TN | AEC line | IAO | | | |
| HLF-133 | Fibroblast | IAO | | | |
| Bronchus Fibroblast | Fibroblast | ENCODE | ENCSR620NSN | ENCFF480YLH | www.encodeproject.org/search/?type=Experiment&assay-term-name=RNA-seq |
| | | "bronchus fibroblast of lung primary cell" | | ENCFF172EXJ | |
| IMR90 | Fibroblast | ENCODE | ENCSR000CTK | ENCFF000HBA | www.encodeproject.org/search/?type=Experiment&assay-term-name=RNA-seq |
| | | "IMR90 female fetal" | | ENCFF000HBI | |
| Fetal Lung | Tissue | ENCODE | ENCSR000AFC | ENCFF002BUL | www.encodeproject.org/search/?type=Experiment&assay-term-name=RNA-seq |
| | | "Lung female fetal tissue (20wks)" | | ENCFF002BUN | |
| AEC-aAT2 | Primary AEC | IAO | | | |
| AEC-fAT2 | Primary AEC | IAO | | | |
| AEC-mAT2 | Primary AEC | Yang et al., 2018 | GSE84273 | GSM2434059 | |
| AEC-aAT1D6 | Primary AEC | IAO | | | |
| AEC-aAT1D2 | Primary AEC | IAO | | | |
| AEC-aAT1D4 | Primary AEC | IAO | | | |
| AEC-fAT1D2 | Primary AEC | IAO | | | |
| AEC-fAT1D4 | Primary AEC | IAO | | | |
| AEC-fAT1D6 | Primary AEC | IAO | | | |
| AEC-mAT1D2 | Primary AEC | Yang et al., 2018 | GSE84273 | GSM2434060 | |
| AEC-mAT1D4 | Primary AEC | Yang et al., 2018 | GSE84273 | GSM2434061 | |
| AEC-mAT1D6 | Primary AEC | Yang et al., 2018 | GSE84273 | GSM2434062 | |
| A427j | LUAD | DBTSS | | DRX015047 | https://trace.ddbj.nig.ac.jp/DRASearch/ |
| A549j | LUAD | DBTSS | | DRX015048 | https://trace.ddbj.nig.ac.jp/DRASearch/ |
| H1299j | LUAD | DBTSS | | DRX015050 | https://trace.ddbj.nig.ac.jp/DRASearch/ |
| H1437j | LUAD | DBTSS | | DRX015051 | https://trace.ddbj.nig.ac.jp/DRASearch/ |
| H1648j | LUAD | DBTSS | | DRX015052 | https://trace.ddbj.nig.ac.jp/DRASearch/ |
| H1650j | LUAD | DBTSS | | DRX015053 | https://trace.ddbj.nig.ac.jp/DRASearch/ |
| H1703j | LUAD | DBTSS | | DRX015054 | https://trace.ddbj.nig.ac.jp/DRASearch/ |
| H1819j | LUAD | DBTSS | | DRX015055 | https://trace.ddbj.nig.ac.jp/DRASearch/ |
| H1975j | LUAD | DBTSS | | DRX015056 | https://trace.ddbj.nig.ac.jp/DRASearch/ |
| H2126j | LUAD | DBTSS | | DRX015057 | https://trace.ddbj.nig.ac.jp/DRASearch/ |
| H2228j | LUAD | DBTSS | | DRX015058 | https://trace.ddbj.nig.ac.jp/DRASearch/ |
| H2347j | LUAD | DBTSS | | DRX015059 | https://trace.ddbj.nig.ac.jp/DRASearch/ |
| H322j | LUAD | DBTSS | | DRX015060 | https://trace.ddbj.nig.ac.jp/DRASearch/ |
| LC2ADj | LUAD | DBTSS | | DRX015062 | https://trace.ddbj.nig.ac.jp/DRASearch/ |
| PC14j | LUAD | DBTSS | | DRX015063 | https://trace.ddbj.nig.ac.jp/DRASearch/ |
| PC3j | LUAD | DBTSS | | DRX015064 | https://trace.ddbj.nig.ac.jp/DRASearch/ |
| PC7j | LUAD | DBTSS | | DRX015065 | https://trace.ddbj.nig.ac.jp/DRASearch/ |
| PC9j | LUAD | DBTSS | | DRX015066 | https://trace.ddbj.nig.ac.jp/DRASearch/ |

IAO = Data generated by authors. Primary AEC sample names were assigned "a", "f", and "m" labels to denote three de-identified lung donors.
ENCODE = Encyclopedia of DNA Elements
DBTSS = Database of Transcription Start Sites (Japan). LUAD cell line sample names were given "j" extensions arbitrarily to distinguish DBTSS data from other sources.
*ENCODE Sample ID name or Gene Expression Omnibus (GEO) record number
**ENCODE data has two accession numbers for paired-end RNA-seq samples. Both samples must be downloaded to get entire dataset
**DBTSS Accession Number will bring you to the webpage where fastq file can be downloaded (Run > DRR > FASTQ)

### Fig 2 – Supplement 1. GO terms associated with Gene Clusters

**A**

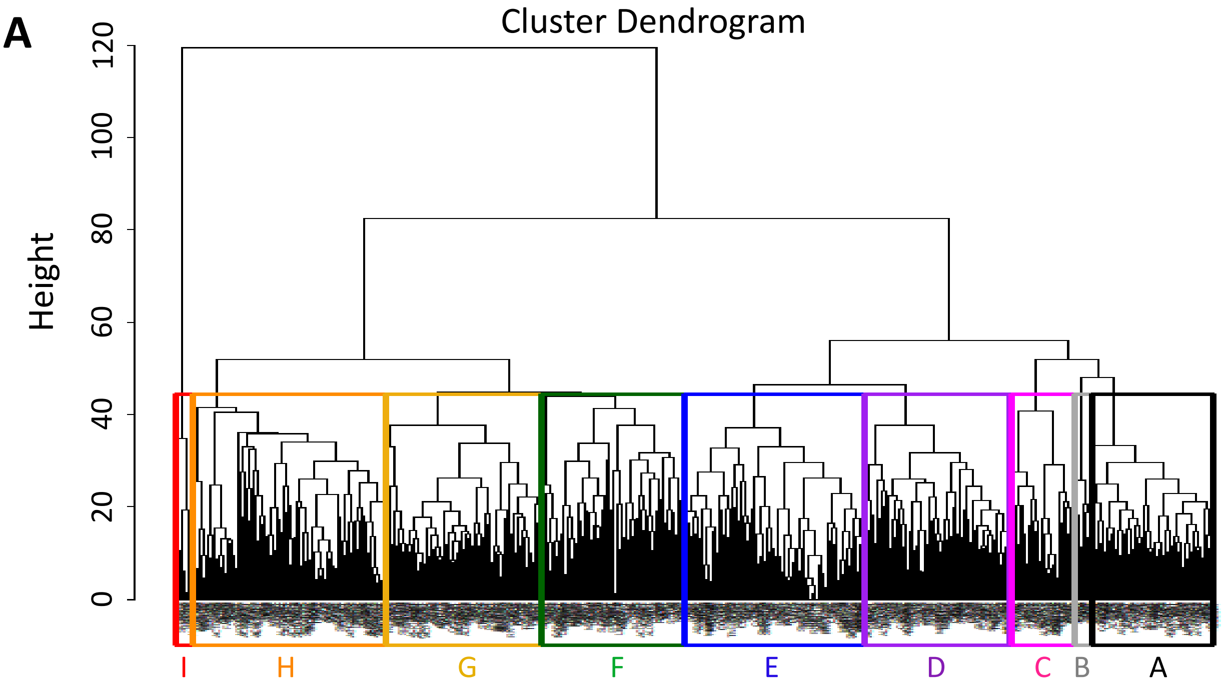


**B**

| Gene Cluster | GO Biological Process | FDR < 0.05 |
|---|---|---|
| A | No statistically significant results | -- |
| B | surfactant homeostasis | 9.36E-08 |
| | respiratory gaseous exchange by respiratory system | 4.46E-08 |
| | positive regulation of phagocytosis | 4.65E-03 |
| | toll-like receptor signaling pathway | 9.17E-03 |
| C | regulation of cell motility | 2.83E-02 |
| | extracellular matrix organization | 1.62E-02 |
| | blood vessel development | 4.45E-02 |
| | tube development | 1.70E-02 |
| D | mesenchymal cell proliferation involved in lung development | 2.16E-02 |
| | morphogenesis of branching epithelium | 3.48E-02 |
| | peptide antigen assembly with MHC class II protein complex | 2.88E-02 |
| | lung lobe morphogenesis | 1.68E-03 |
| | epithelial cell proliferation | 4.27E-03 |
| | regulation of multi-organism process | 3.47E-03 |
| | tube morphogenesis | 4.82E-02 |
| | epithelial tube branching involved in lung morphogenesis | 2.33E-02 |
| | lung epithelium development | 3.08E-02 |
| | cellular response to chemical stimulus | 2.12E-05 |
| | regulation of smooth muscle differentiation | 3.50E-02 |
| E | regulation of cellular response to vascular endothelial growth factor stimulus | 2.24E-02 |
| | cellular response to vitamin | 3.71E-02 |
| | collagen fibril organization | 1.34E-02 |
| | proteoglycan metabolic process | 9.41E-03 |
| | nucleosome assembly | 3..1E-03 |
| | positive regulation of cell-substrate adhesion | 2.20E-02 |
| | animal organ morphogenesis | 1.38E-02 |
| | cellular response to growth factor stimulus | 5.89E-04 |
| | negative regulation of developmental process | 3.69E-02 |
| | negative regulation of multicellular organismal process | 1.96E-02 |
| | tissue development | 4.18E-02 |
| F | No statistically significant results | -- |
| G | tissue development | 2.64E-02 |
| H | No statistically significant results | -- |
| I | No statistically significant results | -- |

**Figure 2 – Supplement 1. GO terms for each gene cluster identified from unsupervised hierarchical clustering. A)** Cluster dendrogram generated using the "Complete" agglomeration method on Euclidean distances to determine gene membership. Colored boxes highlight 9 main gene clusters. **B)** Associated GO terms from the "Biological Processes" annotation data set for each gene cluster determined by the Gene Ontology Consortium's PANTHERv14.1 Overrepresentation Test (Released 20200728) using Fisher's Exact Test with FDR corrected p-values < 0.05. "No statistically significant results" means no significant enrichment in observed genes in the GO category compared to the annotated reference list.

### Fig 2 – Source Data 2. Gene memberships within Clusters A, B, and C

| | | | | | | | | |
|---|---|---|---|---|---|---|---|---|
| **Cluster A** | **PGC** | **SCGB3A1** | **GKN2** | **C4BPA** | SCGB1A1 | SCTR | AGR3 | CTSE |
| | AC008268.1 | ATP13A4 | SCGB3A2 | XIST | SFTA2 | CRTAC1 | RNASE1 | WIF1 |
| | SFTA3 | LINC00891 | LINC00261 | **ALPL** | **LPL** | **EDNRB** | **CLDN18** | **COLEC12** |
| | **BPIFB1** | **CXCL17** | SHE | FAM3B | **C2** | **TRIM71** | CLIC6 | GGT6 |
| | SLC44A4 | BMP3 | BAAT | **IRX1** | **TNIP3** | ARHGEF38 | AL121722.1 | ALOX15B |
| | PIP5K1B | KLHDC7A | VSIG2 | VWA2 | SEC14L6 | **HLA-DOA** | **LRP2** | ADGRD1 |
| | **CNR1** | MXRA5 | **LYZ** | C16orf89 | CHI3L2 | ACADL | C11orf96 | AP003498.2 |
| | **HLA-DPB1** | **COLCA1** | SCNN1G | | | | | |
| **Cluster B** | **SFTPA2** | **AQP4** | **SFTPC** | **SFTPA1** | **NAPSA** | SFTPB | **PIGR** | **SFTPD** |
| **Cluster C** | COL1A2 | SLC34A2 | AL355075.4 | COL6A3 | SLC6A14 | CEACAM6 | **CXCL8** | **GREM1** |
| | SCARNA10 | ITGB6 | SCARNA2 | AL356488.2 | SCARNA7 | CCND2 | **SPARC** | KRT17 |
| | **ADGRF5** | RN7SL3 | **IGFBP5** | **ANPEP** | **MMP2** | **S100A14** | HIST1H1E | SELENOP |
| | **AGR2** | SCARNA13 | ALDH1A1 | TMTC1 | **CD74** | PRSS8 | MPZL2 | |

**Figure 2 – Source Data 2. Gene membership within clusters A, B, and C from unsupervised hierarchical clustering.** Large colored text highlights genes associated with GO terms for "Biological Processes." **Bold black** indicates genes associated with "surfactant homeostasis;" **blue** indicates genes associated with "regulation of immune system process;" **purple** indicates genes associated with "response to external biotic stimuli;" **red** indicates genes associated with "tube development." Gene names with **mixed colors** indicate associations with multiple GO term annotations. GO terms were determined by the Gene Ontology Consortium's PANTHERv14.1 Overrepresentation Test (Released 20200728) using Fisher's Exact Test with FDR corrected p-value cutoff of $p < 0.05$.

### Fig 2 – Source Data 3. List of 75 lung-related genes curated from the literature

| Gene_Symbol | Cell Type marker | Gene_Symbol | Cell Type marker |
|---|---|---|---|
| SFTPC | AT2 | SCGB1A1 | Clara |
| SFTPA1 | AT2 | SCGB3A2 | Clara |
| SFTPA2 | AT2 | CHAD | Clara |
| SFTPB | AT2 | UPK3A | Clara |
| SFTPD | AT2 | NUPR1 | Clara |
| ABCA3 | AT2 | CD200 | Clara |
| LPCAT1 | AT2 | KRT15 | Clara |
| NKX2-1 | AT2 | COL23A1 | Clara |
| CD36 | AT2 | CCND2 | Clara |
| LAMP3 | AT2 | NKX2-5 | Basal cell |
| EGFL6 | AT2 | SOX2 | Basal cell |
| SLC34A2 | AT2 | TP63 | Basal cell |
| DLK1 | AT2 | KRT5 | Basal cell |
| FABP5 | AT2 | KRT14 | Basal cell |
| SOAT1 | AT2 | ITGB4 | Basal cell |
| SCD | AT2 | JAG1 | Ciliated |
| NAPSA | AT2 | LYPD2 | Ciliated |
| ETV5 | AT2 | LRRC23 | Ciliated |
| PDPN | AT1 | CCDC39 | Ciliated |
| AQP5 | AT1 | FOXJ1 | Ciliated |
| AGER | AT1 | STK33 | Ciliated |
| TSPAN8 | AT1 | NCS1 | Ciliated |
| EMP2 | AT1 | CCDC113 | Ciliated |
| DPYSL2 | AT1 | CKAP2L | Ciliated |
| GPRC5A | AT1 | EFHC1 | Ciliated |
| CAV1 | AT1 | EFCAB10 | Ciliated |
| LMO7 | AT1 | NEK10 | Ciliated |
| AKAP5 | AT1 | TEKT4 | Ciliated |
| CLIC5 | AT1 | DTL | Ciliated |
| CLDN18 | AT1 | FAM161A | Ciliated |
| IGFBP6 | AT1 | FHAD1 | Ciliated |
| TIMP3 | AT1 | FANK1 | Ciliated |
| S100A6 | AT1 | HS6ST2 | Ciliated |
| AHNAK | AT1 | DNALI1 | Ciliated |
| COL4A3 | AT1 | KNDC1 | Ciliated |
| HOPX | AT1 | LRRIQ1 | Ciliated |
| | | MCM8 | Ciliated |
| | | CCDC40 | Ciliated |
| | | MELK | Ciliated |

Genes and cell type marker designations were taken from Treutlein et al. (2014) and Xu et al. (2016).

### Fig 2 – Supplement 2. AEC lines express lung-related genes more highly than lung fibroblasts

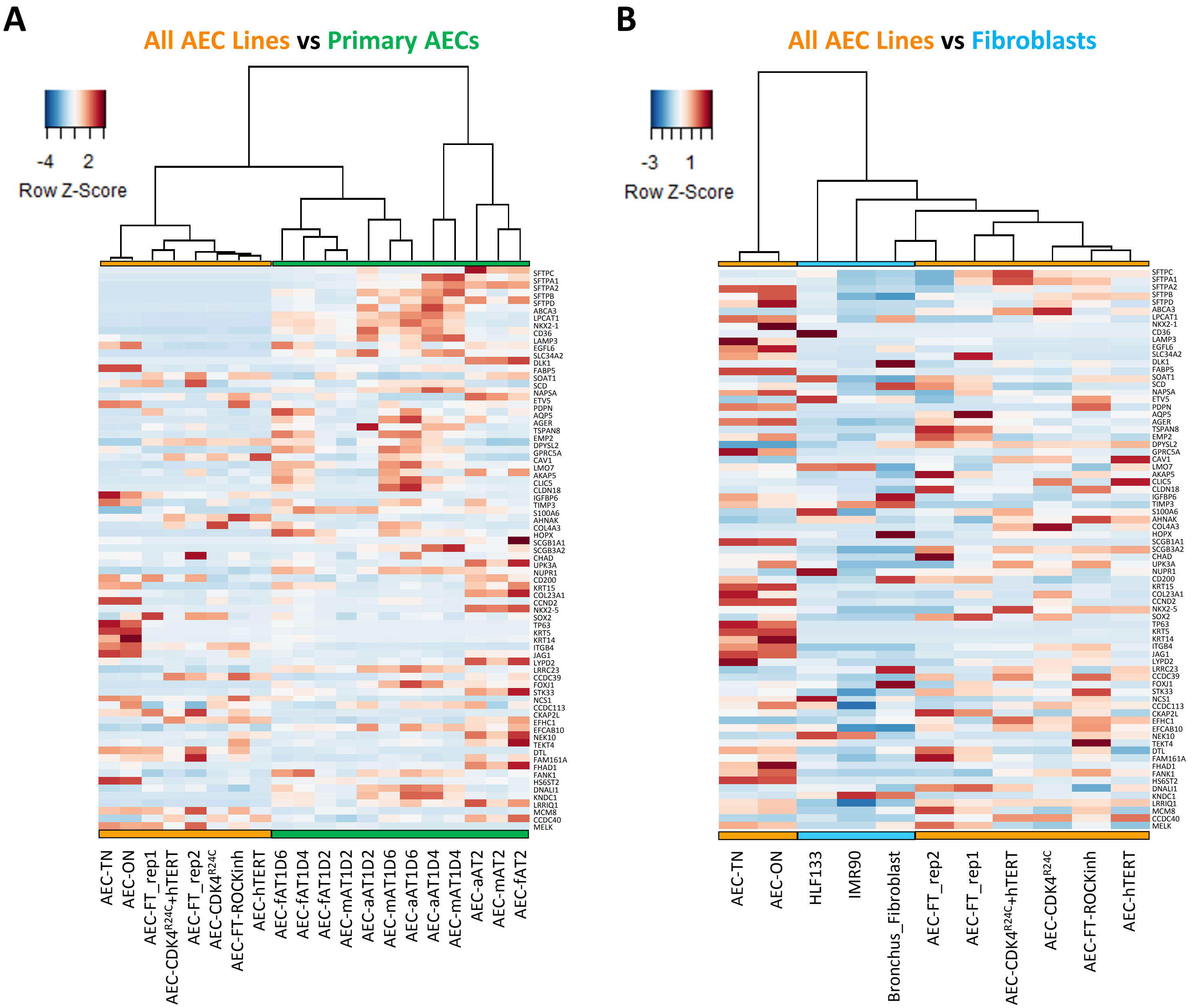


**Figure 2 – Supplement 2. Alveolar epithelial cell lines express lung markers more highly than lung fibroblasts.** FPKM normalized values for all AEC lines, primary AECs, and lung fibroblasts were subsetted on 75 lung-related genes curated from the literature. Unsupervised clustering was performed on sample columns. **A)** Heat map of all AEC lines and primary human AECs. **B)** Heatmap of all AEC lines and lung fibroblasts clustered by Ward's agglomerative method and the Manhattan distance metric. Colored bars indicate sample groups: **orange**, derived AEC lines; **green**, primary AECs (both AT2 and AT1-like); **blue**, lung fibroblasts. Primary AT2 cells are labeled as "—AT2," with "f," "m," and "a" indicating three separate donor lungs. Primary AT1-like cells are labeled as "—D2, " "—D4, "—D6," with numbers representing days transdifferentiated in culture.

**Fig 3 – Supplement 1. AEC lines grown in ROCK inhibitor media express lung progenitor markers, SOX9 and SOX2**

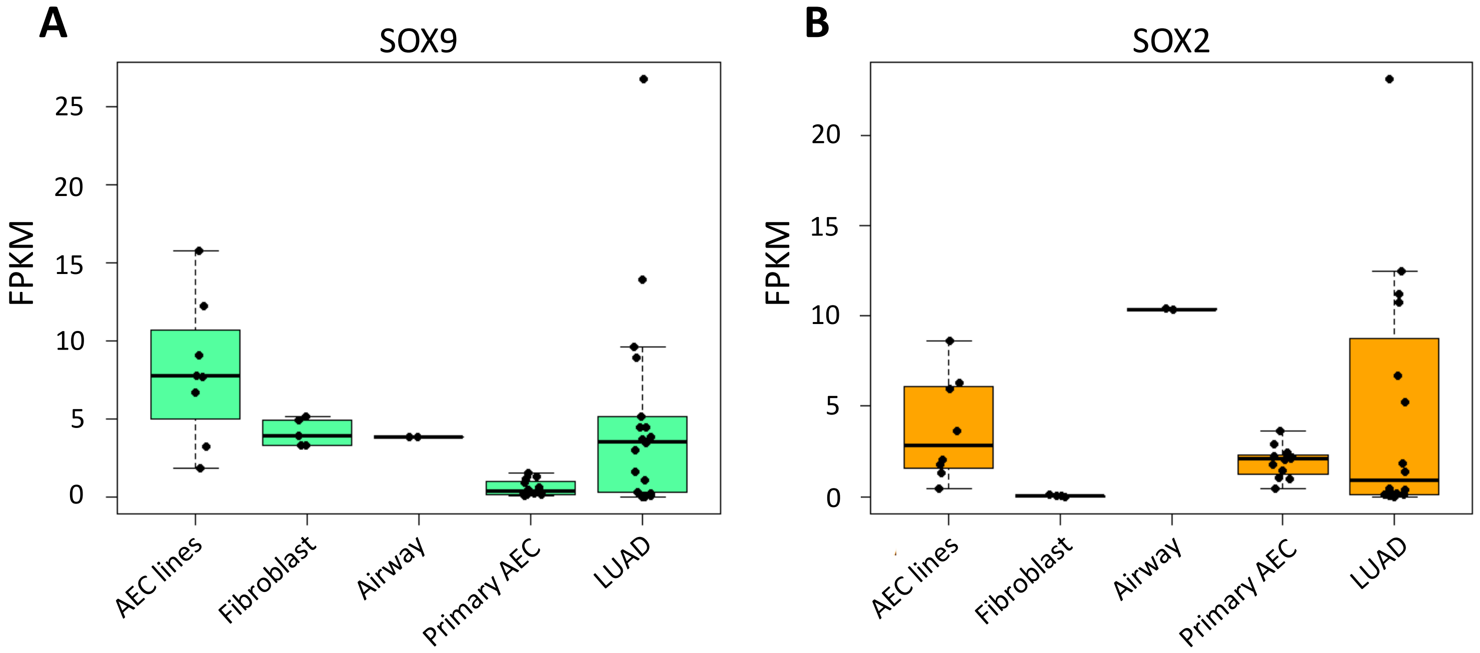


**Figure 3 – Supplement 1. FPKM expression values of SOX9 and SOX2.** Box-and-whiskers plots of normalized FPKM values for **A)** SOX9 and **B)** SOX2 progenitor markers for all AEC lines, lung fibroblasts, airway epithelial cells, primary AECs, and LUAD cancer lines.

### Fig 4 – Supplement 1. A549 lung cancer cells form dense clusters in 3D culture

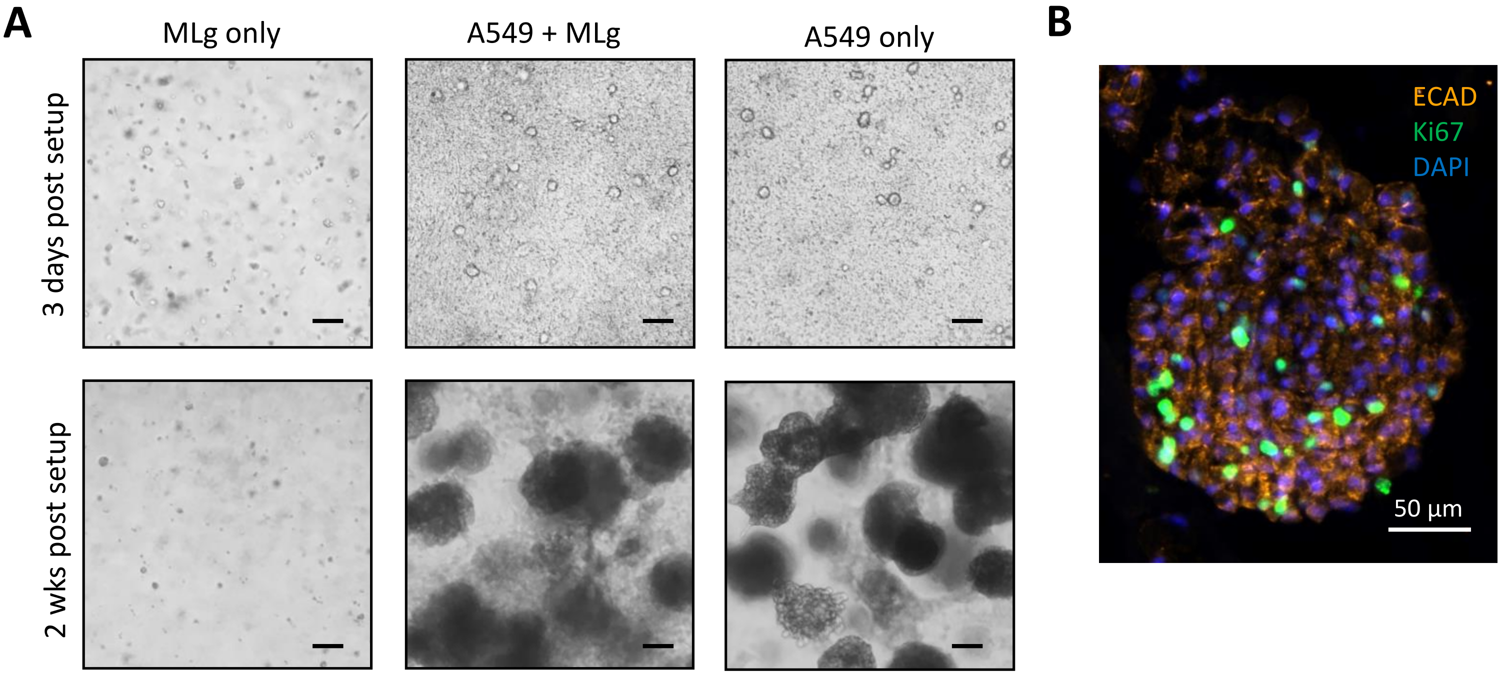


**Figure 4 – Supplement 1. A549 lung adenocarcinoma cells form dense cell clusters in 3D culture. A)** A549 cells, a lung adenocarcinoma cancer cell line derived from a patient lung tumor in 1971, was cultured under 3D organotypic conditions similar to AEC-LgT cell lines. A549s formed dense cell clusters within 2 weeks both in the presence and absence of MLg fibroblasts. Scale, 100 µm. **B)** A549 cell clusters are composed of Ki67$^{+}$ proliferating cells and ECAD$^{+}$ cells. Sectioned A549 cell clusters reveal an absence of structured lumens.

### Fig 4 – Supplement 2. AEC-LgT organoids are not detected in the absence of MLgs

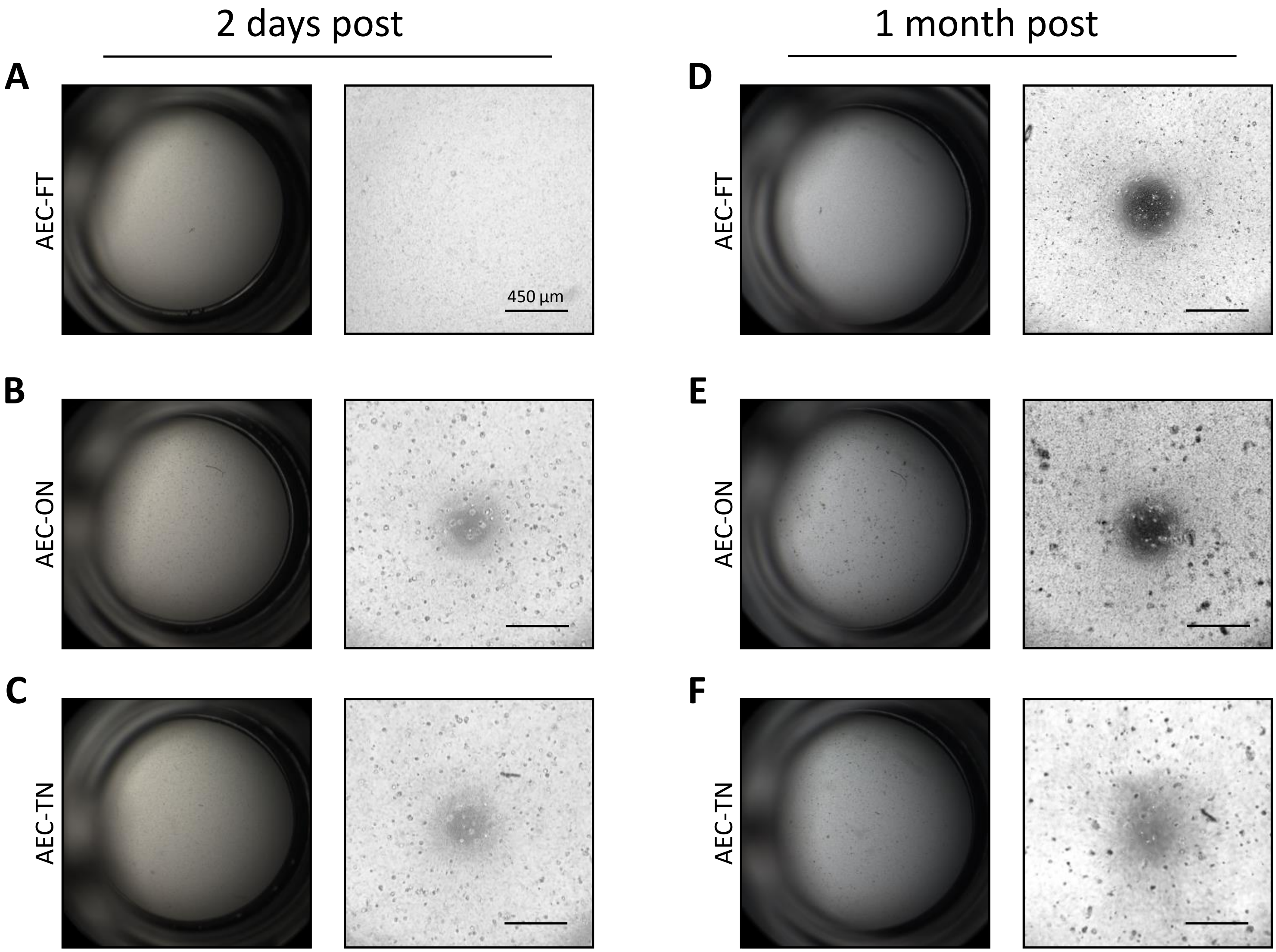


**Figure 4 – Supplement 2. AEC-LgT organoids are not detected when cultured in the absence of MLg fibroblasts.** Representative images of AEC-LgT 3D cultures at 2 days and 1 month post culture set up. **A-C)** Brightfield whole-well images taken at 2.5X magnification (left panels) and zoomed 4X magnification (right panels) after 2 days in culture. Scale bar, 450 µm. **D-F)** Brightfield images taken at 2.5X and 4X magnification of 3D cultures after 1 month of growth. Cellular debris was more noticeable in all wells. Scale bar, 450 µm.

### Fig 4 – Source Data 1. Summary of AEC-LgT organoid growth metrics

| | Organoid size | | | % Efficiency |
|---|---|---|---|---|
| Cell Line | Median (µm) | Mean ± stdev (µm) | Range (µm) | Mean ± stdev (%) |
| AEC-FT | 64 | 82 ± 62 | 25 - 445 | 0.35 ± 0.1 |
| AEC-ON | 108 | 144 ± 106 | 24 - 661 | 1.2 ± 0.5 |
| AEC-TN | 44 | 63 ± 44 | 25 - 427 | 0.2 ± 0.08 |

### Fig 4 – Supplement 3. Additional brightfield images of AEC-LgT organoids to show diversity in shape and size at 2 months

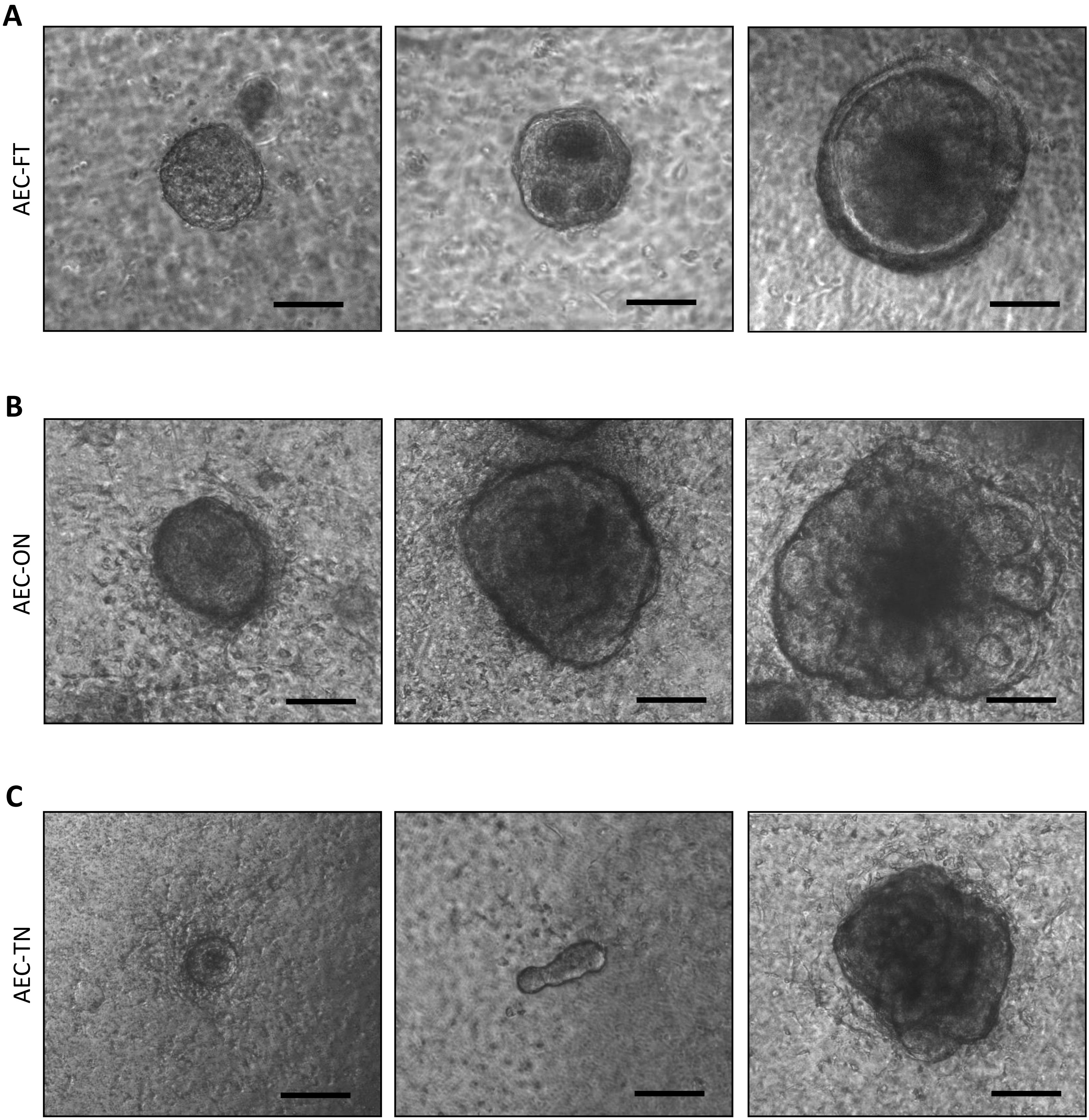


**Figure 4 – Supplement 3. Additional brightfield images of AEC-LgT-derived organoids in 3D co-culture showing diversity in size and shape.** Brightfield images of organoids after 2 months of culture were taken at 4X magnification. Each panel shows a different organoid in an independent well. Scale bar, 100 µm. **A)** AEC-FT, **B)** AEC-ON, and **C)** AEC-TN cell lines.

**Fig 4 – Supplement 4. AEC-LgT organoids express AT1-like markers and not AT2 cell marker, proSPC**

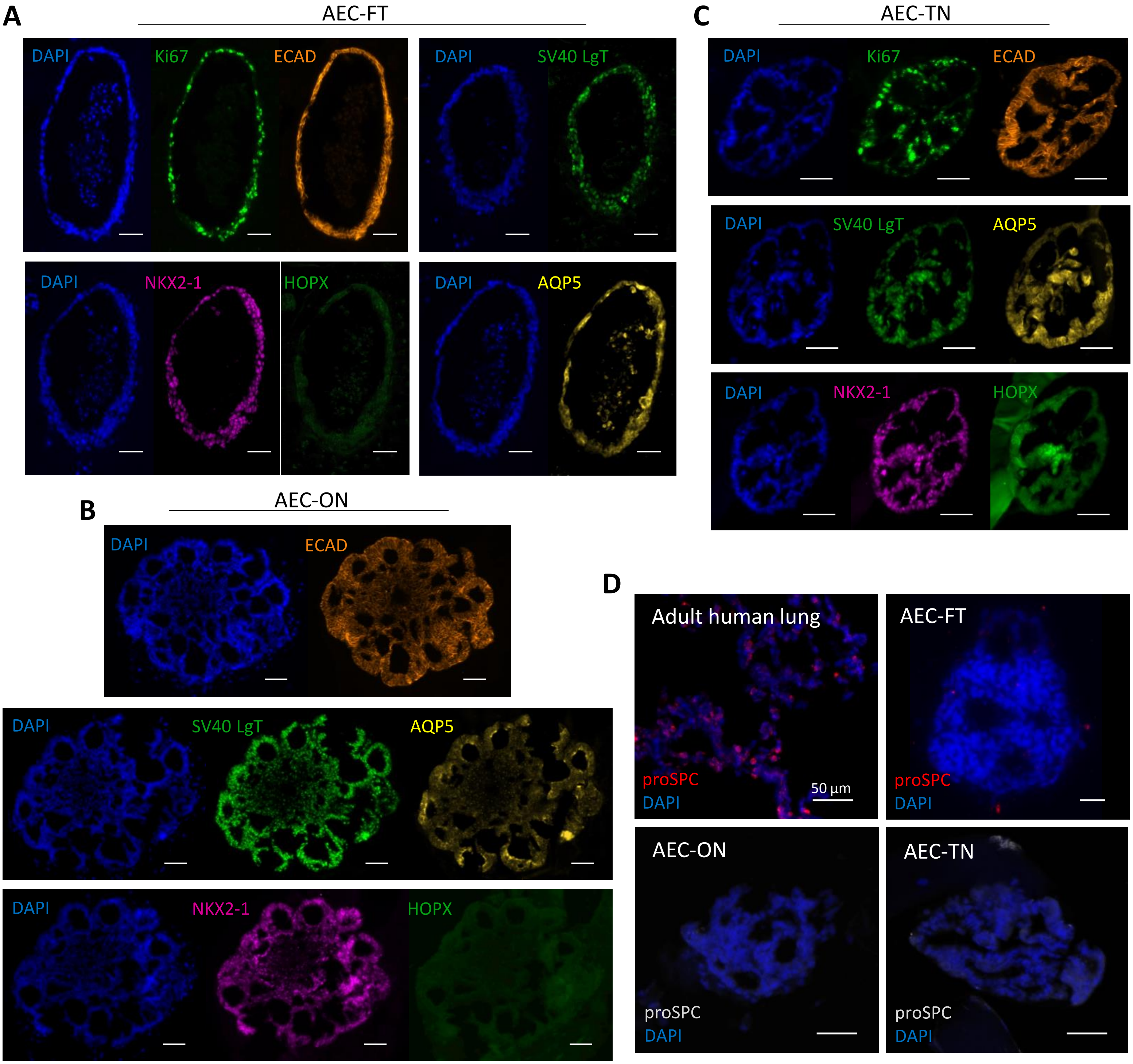


**Figure 4 – Supplemental 4. AEC-LgT-derived organoids express AT1-like markers and not AT2 cell marker, proSPC. A)** AEC-FT, **B)** AEC-ON, and **C)** AEC-TN single-channel fluorescence images for corresponding merged organoid IF stainings shown in Figure 4. Scale bars, 50 µm. **D)** Representative images of stained organoids for AT2 cell marker, proSPC. Adult human lung tissue was used as positive control for proSPC staining. Scale bars, 50 µm.

#### Fig 4 – Supplemental 5. Additional AT2 and AT1 cell marker stainings in 2D and 3D

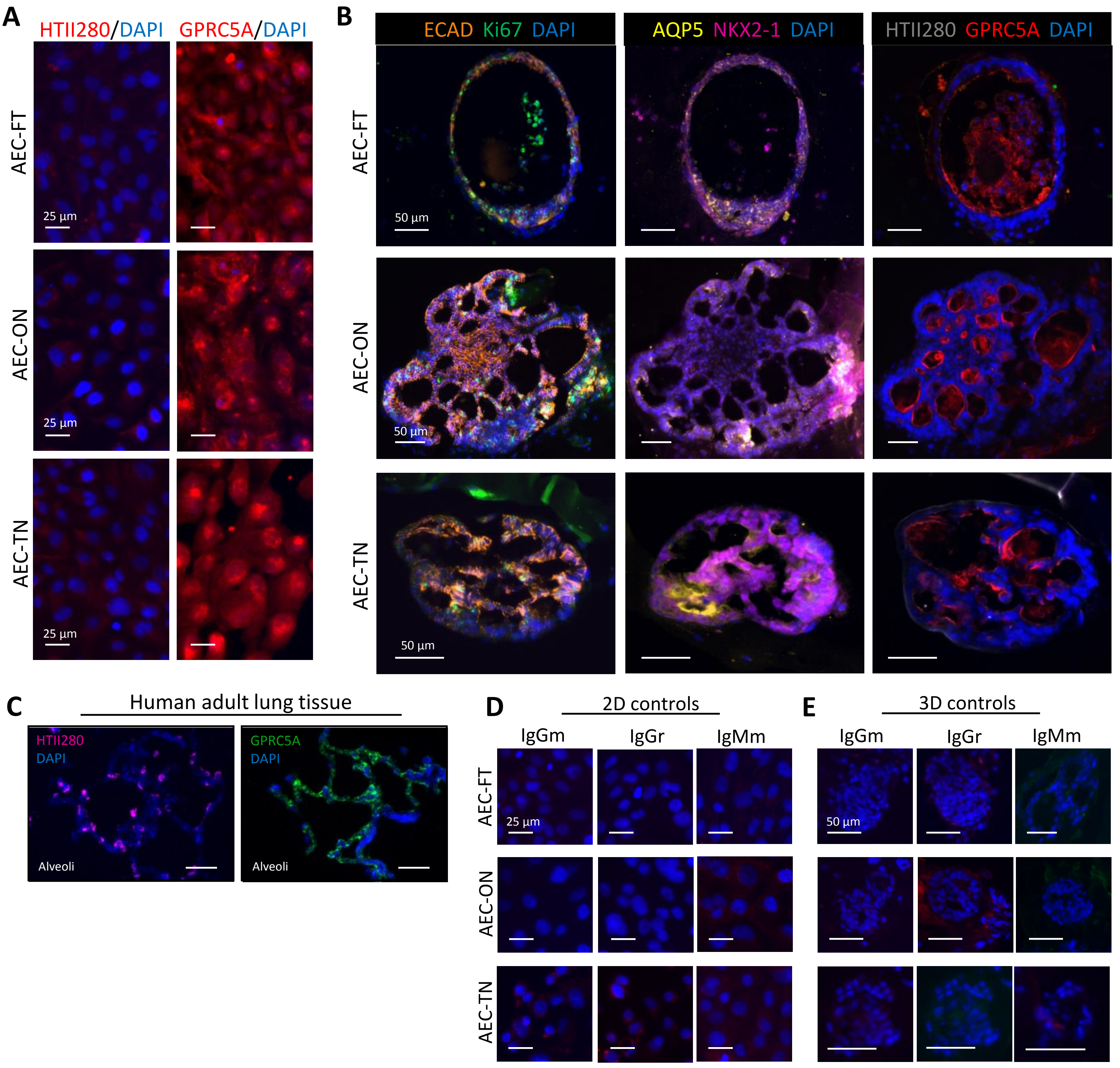


**Figure 4 - Supplement 5. Additional AT2 and AT1 cell marker stainings in 2D and 3D. A)** AEC-FT, AEC-ON, and AEC-TN cells were plated in 2D culture and stained for AT2 cell marker, HTII280, and the recently identified AT1 marker, GPRC5A. Scale bar, 25 µm. **B)** IF staining of AEC-LgT organoid serial sections for the same markers showing subpopulations of cells expressing GPRC5A at the apical membrane. Scale bar, 50 µm. **C)** Human adult lung sections were used as positive controls for antibody specificity. HTII280 and GPRC5A are expressed by alveolar epithelial cells in the distal lung. Scale bar, 50 µm. Negative isotype controls, IgG mouse (IgGm), IgG rabbit (IgGr), and IgM mouse (IgMm) performed on **D)** monolayer cultures and **E)** organoids. For AEC-TN organoids, IgGm and IgGr were probed by double IF staining and shown in their respective fluorescence channels. Scale bar for D), 25 µm. Scale bar for E), 50 µm.

### Fig 6 – Supplemental 1. AEC-FT and AEC-TN organoids respond differently to WNT and FGF pathway activation

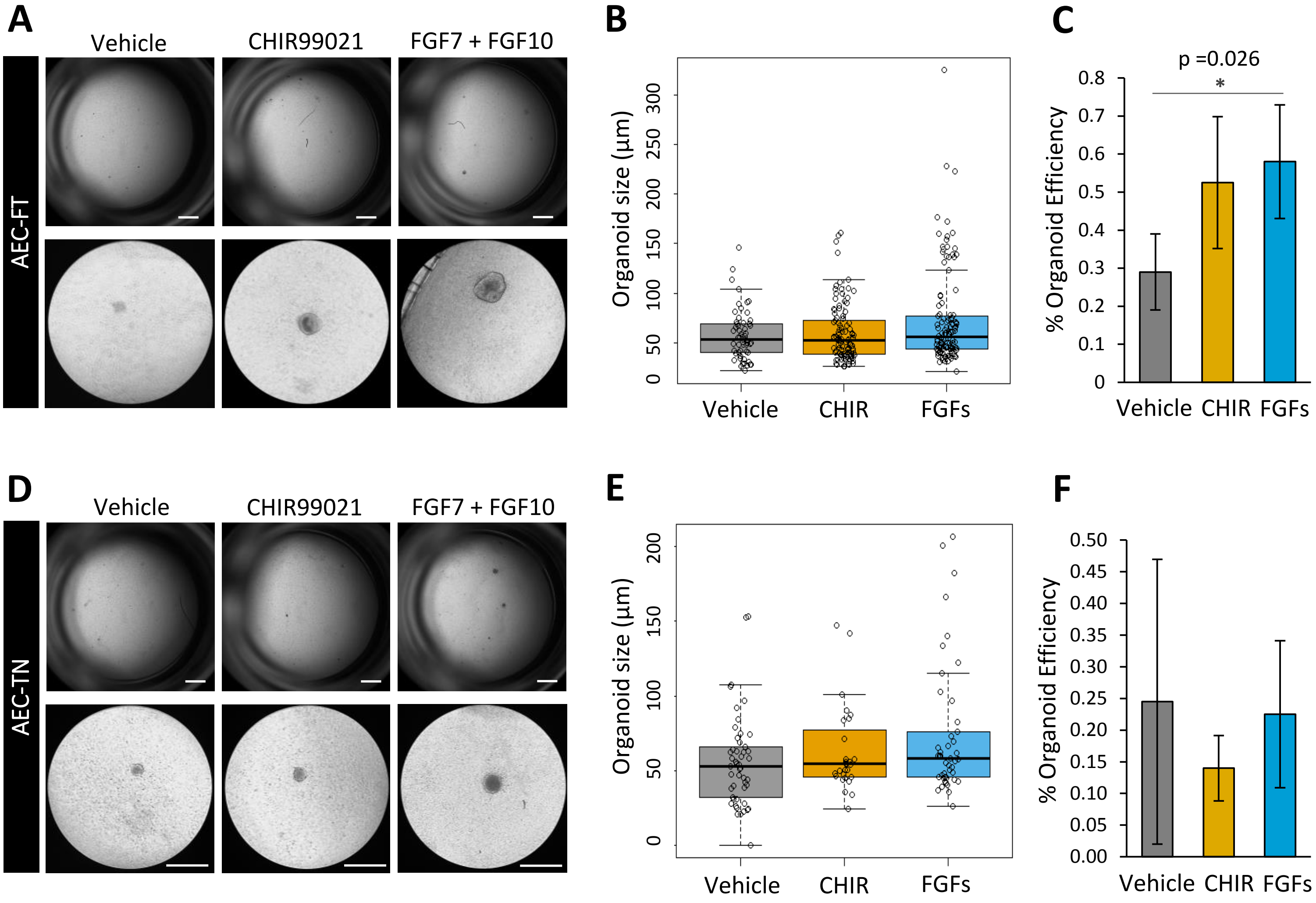


**Figure 6 - Supplement 1. Treatment of AEC-FT and AEC-TN cells in 3D cultures.** AEC-FT and AEC-TN 3D co-cultures were treated with either vehicle (DMSO), 1 µM CHIR99021, or a mix of 10 ng/mL FGF7 and 10 ng/mL FGF10 (FGFs) for 2 months. **A** and **D)** Whole-well and 10X magnification images of a representative well, per treatment condition. Top scale bar, 1000 µm, bottom scale bar 360 µm. **B and E)** Organoid size and **C and F)** organoid formation efficiency were measured for organoids of diameter > 20 µm after 2 months. Plotted values are centered on mean ± standard deviation; n = 4 independent experiments in technical duplicates. *p<0.05 by nonparametric Wilcoxon test.

### Fig 6 – Source Data 1. Summary of growth metrics for treated organoids

| | | Organoid size | | | | % Efficiency | |
|---|---|---|---|---|---|---|---|
| Cell Line | Treatment | Median (µm) | Mean ± stdev (µm) | Range (µm) | p-value* | Mean ± stdev (%) | p-value* |
| AEC-FT | vehicle (DMSO) | 53 | 57 ± 25 | 21 - 146 | -- | 0.3 ± 0.1 | -- |
| | CHIR99021 | 53 | 60 ± 29 | 26 - 158 | 0.83 | 0.5 ± 0.2 | 0.18 |
| | FGF7+FGF10 | 56 | 73 ± 49 | 21 - 172 | 0.12 | 0.6 ± 0.2 | 0.026 |
| AEC-ON | vehicle (DMSO) | 83 | 126 ± 12 | 21 - 613 | -- | 1.8 ± 0.4 | -- |
| | CHIR99021 | 155 | 207 ± 29 | 24 - 659 | $< 2.2 \times 10^{-16}$ | 1.2 ± 0.4 | 0.11 |
| | FGF7+FGF10 | 174 | 202 ± 27 | 30 - 813 | $< 2.2 \times 10^{-16}$ | 1.6 ± 0.3 | 0.88 |
| AEC-TN | vehicle (DMSO) | 53 | 56 ± 10 | 20 - 153 | -- | 0.2 ± 0.2 | -- |
| | CHIR99021 | 54 | 64 ± 12 | 24 - 142 | 0.35 | 0.1 ± 0.05 | 0.77 |
| | FGF7+FGF10 | 58 | 76 ± 9 | 26 - 207 | 0.052 | 0.2 ± 0.1 | 0.77 |

* p-value calculated by nonparametric Wilcoxon test
